# Intrinsic Mechanisms in the Gating of Resurgent Na^+^ Currents

**DOI:** 10.1101/2021.04.09.439048

**Authors:** Joseph L. Ransdell, Jonathan D. Moreno, Druv Bhagavan, Jonathan R. Silva, Jeanne M. Nerbonne

## Abstract

The resurgent component of the voltage-gated sodium current (I_NaR_) is a depolarizing conductance, revealed on membrane hyperpolarizations following brief depolarizing voltage steps, which has been shown to contribute to regulating the firing properties of numerous neuronal cell types throughout the central and peripheral nervous systems. Although mediated by the same voltage-gated sodium (Nav) channels that underlie the transient and persistent Nav current components, the gating mechanisms that contribute to the generation of I_NaR_ remain unclear. Here, we characterized Nav currents in mouse cerebellar Purkinje neurons, and used tailored voltage-clamp protocols to define how the voltage and the duration of the initial membrane depolarization affect the amplitudes and kinetics of I_NaR_. Using the acquired voltage-clamp data, we developed a novel Markov kinetic state model with parallel (fast and slow) inactivation pathways and, we show that this model reproduces the properties of the resurgent, as well as the transient and persistent, Nav currents recorded in (mouse) cerebellar Purkinje neurons. Based on the acquired experimental data and the simulations, we propose that resurgent Na^+^ influx occurs as a result of fast inactivating Nav channels transitioning into an open/conducting state on membrane hyperpolarization, and that the decay of I_NaR_ reflects the slow accumulation of recovered/opened Nav channels into a second, alternative and more slowly populated, inactivated state. Additional simulations reveal that extrinsic factors that affect the kinetics of fast or slow Nav channel inactivation and/or impact the relative distribution of Nav channels in the fast- and slow-inactivated states, such as the accessory Navβ4 channel subunit, can modulate the amplitude of I_NaR_.

**SUMMARY:** The resurgent component of the voltage-gated sodium current (I_NaR_) is revealed on membrane *hyperpolarizations* following brief depolarizing voltage steps that activate the rapidly activating and inactivating, transient Nav current (I_NaT_). To probe the mechanisms contributing to the generation and properties of I_NaR_, we combined whole-cell voltage-clamp recordings from mouse cerebellar Purkinje neurons with computational modeling to develop a novel, blocking particle-independent, model for the gating of I_NaR_ that involves two parallel inactivation pathways, and we show that this model recapitulates the detailed biophysical properties of I_NaR_ measured in mouse cerebellar Purkinje neurons.

## INTRODUCTION

Voltage-gated sodium (Nav) channels open rapidly on membrane depolarization and underlie the generation of action potentials in many excitable cells, including skeletal and cardiac muscle, as well as central and peripheral neurons. The pore-forming (α) subunits of Nav channels, Nav1.1 to Nav1.9, belong to the “S4” superfamily of voltage-gated ion channel genes (Catterall, 2010). Each Nav α subunit comprises four homologous domains (DI-DIV) with six transmembrane spanning segments (S1-S6) (Noda et al., 1984). The S1-S4 segments in each domain form the four voltage sensing domains (VSDs) that activate on membrane depolarization (Noda et al., 1984, Guy and Seetharamulu, 1986), resulting in channel opening and Na^+^ influx. Nav channels conduct Na^+^ when the VSDs of domains I, II and III move outwardly to an activated conformation (Bezanilla, 2000). Following opening, fast inactivation occurs (Bezanilla and Armstrong, 1977), mediated by a hydrophobic (IFM) motif in the cytosolic DIII–DIV linker that binds to a site near the pore that is revealed on activation of the DIII and DIV VSDs (Bosmans et al., 2008, Capes et al., 2013, Hsu et al., 2017, Cha et al., 1999). Thus, fast inactivation of open Nav channels occurs when all four VSDs are in the outward/activated position (Capes et al., 2013). If domains I, II and III are activated and DIV is in the deactivated position, however, the IFM motif does not bind, which results in the generation of a non-inactivating or persistent Nav current (I_NaP_) component (Chanda and Bezanilla, 2002, Horn et al., 2000).

An additional Nav current component, that is observed on membrane *hyperpolarizations* following brief depolarizing voltage steps and referred to as the resurgent component of Nav current (I_NaR_), was first described in isolated, postnatal day 8-14 rat cerebellar Purkinje neurons (Raman and Bean, 1997). Although linked to the regulation of the spontaneous firing of action potentials in cerebellar neurons (Raman and Bean, 1997, 1999), I_NaR_ was subsequently identified in more than twenty types of neurons in the central and peripheral nervous systems, only some of which are spontaneously active (Lewis and Raman, 2014) suggesting that I_NaR_ likely plays diverse functional roles in regulating neuronal excitability. Although flowing through the same Nav channels as I_NaT_ and I_NaP_, the time- and voltage-dependent properties of I_NaR_ are distinct (Lewis and Raman, 2014). In addition to being revealed on membrane *hyperpolarizations* presented after brief (∼5 ms) depolarizing voltage steps that evoke I_NaT_ (Khaliq et al., 2003), for example, the time courses of I_NaR_ activation and decay are much slower than I_NaT_. These experimental observations were interpreted as suggesting a Nav channel gating model with two distinct mechanisms contributing to inactivation: a conventional, fast inactivation mechanism in which channels recover from inactivation without passing through an open conducting state; and, a second mechanism, favored by brief depolarizations, in which channels recover from inactivation by passing through an open, conducting state (Raman and Bean, 2001). It was further suggested that the second mechanism was consistent with a voltage-dependent process whereby Nav cannels, opened on depolarization, are blocked by an endogenous ‘blocking’ particle that occludes the pore, driving channels into an “open-blocked” state (Raman and Bean, 2001). On subsequent membrane hyperpolarization, the blocking particle is displaced, and Na^+^ flows through unblocked/open Nav channels, generating the resurgent Nav current (Raman and Bean, 2001). Importantly, in this gating scheme, blocked Nav channels do not inactivate and inactivated channels are not blocked, i.e., a Nav channel cannot be blocked and inactivated simultaneously (Lewis and Raman, 2014).

Studies focused on defining the molecular mechanism(s) underlying the generation of I_NaR_ revealed that proteases (e.g., trypsin/chymotrypsin) that act at positively charged and aromatic/hydrophobic amino acid residues eliminate I_NaR_, while increasing I_NaT_, observations interpreted as supporting the blocking particle model and suggesting that the putative blocker was a protein within the Nav channel complex (Grieco et al., 2002). Attention focused quickly on the transmembrane accessory Navβ4 subunit, which has a short cytosolic tail with several positive charges and multiple aromatic/hydrophobic residues (Grieco et al., 2005). Clear support for a role for Navβ4 was provided in experiments in which intracellular application of a synthetic Navβ4 peptide containing the tail sequence (β4154-167), following elimination of the resurgent Nav current by trypsin or chymotrypsin, rescued I_NaR_ in isolated cerebellar Purkinje neurons (Grieco et al., 2005). In addition, experiments on CA3 pyramidal neurons, which lack I_NaR_ and Navβ4, demonstrated that the application of the β4154-167 peptide generated resurgent Nav currents (Grieco et al., 2005). Further support for a critical role for Navβ4 was provided with the demonstration that treatment of mouse cerebellar granule neurons with small interfering RNAs (siRNAs) targeting *Scn4b* (Navβ4) resulted in the loss of I_NaR_ and that the subsequent exposure of *Scn4b*-siRNA-treated cells to the β4154-167 peptide rescued I_NaR_ (Grieco et al., 2005, Bant and Raman, 2010).

In experiments designed to test directly the hypothesis that Navβ4 is *required* for the generation of I_NaR_, however, we found that I_NaR_ was reduced (by ∼ 50%), but *not* eliminated, in cerebellar Purkinje neurons in (*Scn4b^-/-^*) mice lacking Navβ4 (Ransdell et al., 2017). Similar results were subsequently obtained in experiments conducted on cerebellar Purkinje neurons isolated from another *Scn4b^-/-^* mouse line (White et al. (2019). In addition, it has been reported that I_NaR_ is readily measured in *Scn4b^-/-^* striatal neurons (Miyake et al., 2014). These observations clearly suggest that additional mechanisms contribute to the generation of I_NaR_. One possibility is that there are additional open channel blocking molecules expressed in Purkinje (and other) neurons. Support for this hypothesis was provided in studies showing that I_NaR_ is decreased (but, again, not eliminated) in neonatal mouse cerebellar Purkinje isolated from animals harboring a targeted disruption in the *Fgf14* (which encodes intracellular fibroblast growth factor 14, iFGF14) locus (White et al., 2019), as well as in wild type neonatal Purkinje cells following exposure to an interfering RNA targeting the *Fgf14b* variant (Yan et al., 2014). Interestingly, it was also reported that I_NaR_ was readily detected in Purkinje neurons isolated from neonatal animals lacking both *Scn4b* and *Fgf14* (White et al. 2019). It is certainly possible and that there are additional, yet to be discovered, endogenous open channel blockers that contribute to the generation of I_NaR_. Alternatively, it seemed possible to us that there is an intrinsic gating mechanism(s) by which Nav channels can produce resurgent current. Here, we present the results of experimental and modeling efforts designed to explore the latter hypothesis directly, and we provide evidence for a “blocking-particle independent” mechanism in the generation of I_NaR_ in cerebellar Purkinje neurons.

## METHODS

### Animals

All animal experiments were performed in accordance with the guidelines published in the National Institutes of Health Guide for the Care and Use of Laboratory Animals. Protocols were approved by the Washington University Institutional Animal Care and Use Committee (IACUC). Postnatal day 12-16 (P12-P16) male and female C57BL6/J (Jackson laboratories) mice were used in the experiments reported here.

### Isolation of neonatal cerebellar Purkinje neurons

For the preparation of isolated cerebellar Purkinje neurons, postnatal day 12-16 (P12-P16) animals were anesthetized with 1.25% Avertin and brains were rapidly removed and placed in ice-cold isolation medium containing (in mM): 82 Na_2_SO_4_, 30 K_2_SO_4_, 5 MgCl_2_, 10 HEPES, 10 glucose, and 0.001% phenol red (at pH 7.4). Using a scalpel, the cerebellum was removed, minced into small chunks and incubated in isolation medium containing 3 mg/ml protease XXIV at 33°C for 10-15 min. Following this incubation period, the tissue pieces were washed with enzyme-free isolation medium containing 1 mg/ml bovine serum albumin and 1 mg/ml trypsin inhibitor. The tissue pieces were transferred to oxygenated artificial cerebral spinal fluid (ACSF) containing (in mM): 125 NaCl, 2.5 KCl, 1.25 NaH_2_PO_4_, 25 NaHCO_3_, 2 CaCl_2_, 1 MgCl_2_, and 25 dextrose (∼310 mosmol/L) at 22–23°C. Tissue pieces were triturated with a fire-polished glass pipette. An aliquot of the cell suspension was placed on a coverslip in the recording chamber and superfused with fresh ACSF (at a rate of 0.5 ml/min), saturated with 95% O_2_/5% CO_2_, for 25 minutes before beginning electrophysiological experiments.

### Electrophysiological recordings and analysis

Whole-cell voltage-clamp recordings were obtained at room temperature from visually identified cerebellar Purkinje neurons using differential interference contrast optics. Data were collected using a Multiclamp 700B patch clamp amplifier interfaced to a Dell PC with a Digidata 1332 and pCLAMP 10 software (Axon Instruments, Union City, CA, USA). In all recordings, tip potentials were zeroed prior to forming a giga-ohm membrane–pipette seal. Pipette capacitances were compensated using the pCLAMP software. Signals were acquired at 50-100 kHz and filtered at 10 kHz prior to digitization and storage.

For recordings of I_NaR_, the normal ACSF perfusing the cells was switched to ACSF that also contained 5 mM tetraethylammonium chloride (TEACl) and 250μM cadmium chloride (CdCl_2_). To decrease Nav currents and improve the spatial control of the membrane voltage (space-clamp) and enable reliable recordings of I_NaT_ or I_NaP_ (**Figure 6**), ACSF containing (in mM): 25 NaCl, 100 TEACl, 2.5 KCl, 1.25 NaH_2_PO_4_, 25 NaHCO_3_, 2 CaCl_2_, 1 MgCl_2_, 25 dextrose and 0.25 CdCl_2_ was used. Recording pipettes contained (in mM): 110 CsCl, 15 TEACl, 5 4AP, 1 CaCl_2_, 2 MgCl_2_, 10 EGTA, 4 Na_2_-ATP and 10 HEPES, pH 7.25 ∼300 mosmol/L. Alternative ACSF and internal solutions were used in experiments presented in **Figure 2**; these solutions are described in the Results section and in the legend to **Figure 2**. Recording pipettes had resistances of 2-3 MΩ. Series resistances were routinely compensated ≥ 80%. Voltage errors resulting from the uncompensated resistances were always ≤ 4mV and were not corrected. Peak transient Nav conductances (at each test potential) were calculated (using E_reversal_ = +75 mV) in each cell and normalized to the maximal peak transient Nav conductance determined in the same cell. Mean ± SEM normalized peak transient Nav conductances were calculated and plotted as a function of the test potential. The mean ± SEM peak transient Nav conductance activation plots were fitted with a single (equation 1) Boltzmann function shown below:

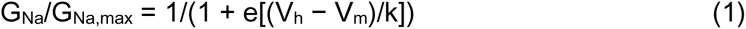

Where V_h_ is the membrane potential of half-maximal activation and k is the slope factor.

To determine the voltage-dependence of steady-state inactivation of the transient Nav currents, a two step protocol was used. From a holding potential of -70 mV, brief (20 ms) voltage steps to various conditioning voltages, between -120 mV and -35 mV, were presented prior to depolarizations to 0 mV to evoke I_NaT_. Peak I_NaT_ amplitudes at 0 mV, evoked from each conditioning potential in each cell, were measured and normalized to the peak I_NaT_ amplitude evoked (at 0 mV) from the -120 mV conditioning step in the same cell. Mean ± SEM normalized peak I_NaT_ amplitudes were plotted as a function of the conditioning voltage and fitted with a single Boltzmann function (equation 2).

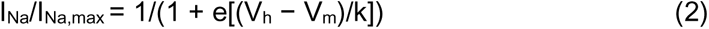

Where V_h_ is the membrane potential of half-maximal inactivation and k is the slope factor. The time constants (tau) of I_NaR_ inactivation were determined from first-order fits to the decay phases of the inward currents evoked on hyperpolarizations to various potentials following brief (5 ms) depolarizing voltage steps to 0 mV.

Data were analyzed using ClampFit (Molecular Devices), Matlab (Mathworks), Microsoft Excel and Prism (GraphPad Software Inc.).

### Simulations using the Raman-Bean model

The Raman-Bean model of Nav channel gating (*Raman and Bean, 2001)* was coded in Matlab using the equations and schematic of the Markov model structure presented in **Figure 7** of Raman and Bean (2001); no changes were made to the published model or parameters (*Raman and Bean, 2001)*. The model simulations were run as described below using the matrix exponential technique (Teed and Silva, 2016).

### Development of a computational model of Nav channel gating

A Markov kinetc state model of Nav channel gating in mouse cerebellar Purkinje neurons was formulated based on our acquired experimental data that led to a hypothesized structure of channel gating states (see: **Figure 3A**). The rate constants in the model were all single exponentials derived from the acquired experimental data and were optimized numerically using described methods (Moreno et al., 2016, Teed and Silva, 2016). All simulations, numerical optimization and data visualization were done in MATLAB 2017B™. Detailed methods are below. Source code is available upon request. The equations were verified by one of the authors (DB)who was not involved in the creation of the model.

The Nav channel gating model developed (see: **Figure 3**) consists of 10 states: 3 closed states (C1, C2, C3), a single open state (O), 2 closed-inactivated states (IC1, IC2), two inactivated states, resulting from fast channel inactivation (IF1, IF2), and an additional, slowly populated inactivated state (IS). The model accurately simulates the experimentally observed kinetics and voltage-dependences of Nav channel gating, including activation, inactivation (closed and open state), deactivation, recovery from fast inactivation, as well as the voltage-dependence of activation and the proportion (relative to I_NaT_) of I_NaR_ and I_NaP_.

### Numerical optimization procedure

All computations were done in MATLAB 2017B™. We coded all voltage protocols and used the matrix exponential technique, which is described in Teed and Silva (2016), for simulations. A modified Nelder Mead Simplex method that allows for constrained optimization (only positive rate constants) was used for simultaneous optimization of the protocols listed below. A cost function for each protocol was defined as the sum of squared differences between the experiments and the simulations. The total cost function (sum of the individual protocol errors) was then minimized and converged when a tolerance of 0.01 for the change of the cost function and 0.01 for the change in parameters was achieved. For further details about this numerical optimization method, see Moreno et al. (2016).

The protocols were as follows:

- ***Steady-state availability:*** For each voltage between -120 mV and -10 mV, the steady-state probabilities of the channel were found. The channel was then depolarized to 0 mV, and the open-state probability was determined. The value of the open state probability was then normalized to the open state probability at -120 mV.
- ***Steady-state activation:*** Channel steady-state was found at -80 mV. The channel was then depolarized to voltages between -77 and 0 mV (in 3 mV increments). For each voltage, the maximum open probability of the channel was calculated, and the conductance, G_Na_, at each voltage was determined. The calculated values were then normalized to G_Na_ at 0 mV.
- ***Tau of deactivation:*** Channel steady-state was found at a holding potential of -90 mV. From steady-state, the channel was depolarizedto voltages between -50 mV and 0 mV (in 5 mV increment), and the time constant (tau) corresponding to 64% decay (1/e) of the peak current was calculated for each voltage.
- ***Recovery from fast inactivation:*** From a steady-state of -90 mV, the channel was depolarized to 0 mV, and the peak current determined. The channel was then allowed to recover at -90 mV for variable time intervals, before being depolarized again to 0 mV. The peak currents during the depolarizations to 0 mV following the various recovery times were determined and normalized to the initial peak current.
- ***Persistent component of the Nav current:*** From a steady-state of -90 mV, the channel was depolarized to 0 mV for 5 ms, and subsequently hyperpolarized to -45 mV for 100 ms. The persistent current recorded after the 100 ms hyperpolarizing step was measured and normalized to the initial peak current.
- ***Voltage-dependence of the ratio of the peak resurgent to peak transient Nav current amplitude***: From a steady-state of -90 mV, the channel was depolarized to 0 mV for 5 ms, and subsequently hyperpolarized to potentials between -5 mV and -80 mV (in 5 mV increments). The peak resurgent current at each hyperpolarized voltage step was determined and normalized to the peak transient inward current evoked at at 0 mV.
- ***Dependence of the peak I_NaR_ amplitude on the duration of the depolarizing voltage step duration:*** From a steady-state of -90 mV, the channel was depolarized to +20 mV for varying times (2 ms to 36 ms) prior to hyperpolarization to -45 mV. The amplitude of I_NaR_ at -45 mV following each depolarizing voltage step to +20 mV (of varying durations) was measured and normalized to the peak I_NaR_ evoked following the 2 ms depolarizing voltage step to _20 mV.
- ***Tau of inactivation of the resurgent Nav current:*** From a steady-state of -90 mV, the channel was depolarized to 0 mV for 5 ms, and subsequently hyperpolarized to potentials ranging from -5 to -45 mV (in 5 mV increments). The time constant (tau) of the exponential decay (1/e) of the resurgent current at each hyperpolarized test potential was determined.
- ***Voltage independence of peak resurgent current:*** Consistent with the experimental data presented in **Figures 2D and 2E**, the peak resurgent current was independent of the potential of the depolarizing voltage step. From a steady-state of -90 mV, the channel was depolarized to membrane potentials ranging from -5 mV to -35 mV (in 5 mV increments) prior to a hyperpolarizing voltage step to -45 mV. The difference between the peak I_NaR_ measured at -45 mV from each depolarized potential was determined, and the mean resurgent peak current determined across all depolarized potentials was minimized. This ensured the peak resurgent current was constant without specifying the magnitude of the resurgent peak *a priori*.

### Optimization of the Scn4b^-/-^ Nav current model

To develop the model for Nav channels lacking *Scn4b*, the optimization routine was restarted from the optimized wild type rate constants (initial conditions). The “***Voltage-dependence of the ratio of the peak resurgent to peak transient Nav current amplitude***” protocol described above was fitted to the experimental data acquired from isolated cerebellar Purkinje neurons from (*Scn4b*^-/-^) animals harboring a targeted disruption in the *Scn4b* locus (Ransdell et al., 2017); the data are presented in **Figure 9A**. All of the other protocols used in the optimization procedure for *Scn4b^-/-^* were the same as those described above for the wild type Nav channel to ensure no other changes to the model.

### Model parameters

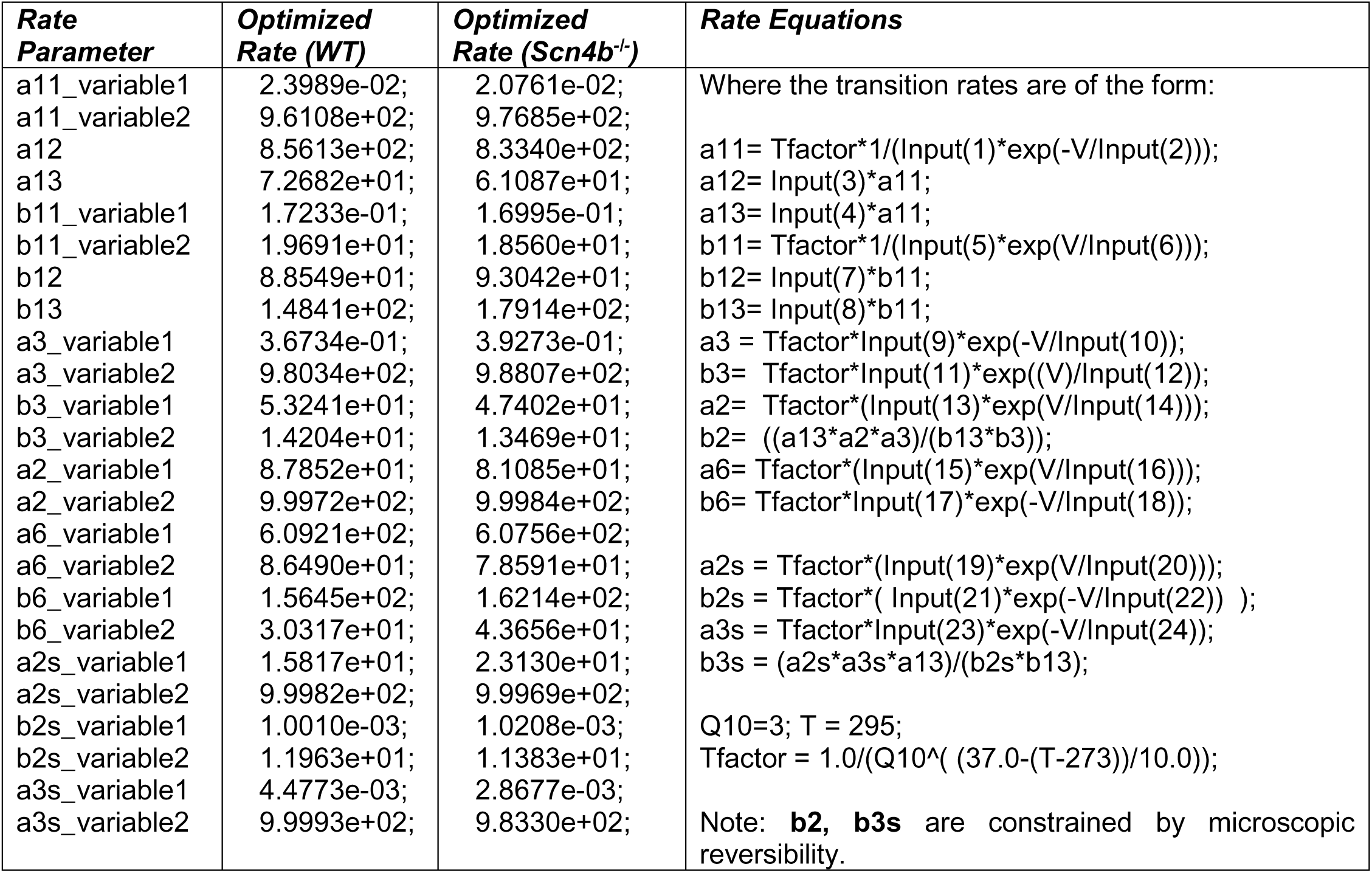

Of note, in the MATLAB script, the Nav channel model developed here contains 24 parameters; these are inputted as a matrix “Input”. For example, Input(1) corresponds to a11_variable1, and Input(2) corresponds to a11_variable2, Input(3) corresponds to a12, and Input(4) corresponds to a13, etc. etc. The transition rate constants are of the form denoted in the right hand column of the table above.

## RESULTS

### The amplitude of I_NaR_ depends on the duration, but not the voltage, of the prior membrane depolarization

In isolated neonatal (P12-P16) mouse cerebellar Purkinje neurons, the fast transient (I_NaT_), persistent (I_NaP_) and resurgent (I_NaR_) Nav current components can be distinguished using voltage-clamp protocols that take advantage of the unique time- and voltage-dependent properties of the three current components (**Figure 1A**). On membrane depolarization, for example, I_NaT_ activates fast and subsequently decays rapidly to a steady-state (persistent) level of inward current, I_NaP_ (**Figure 1A**). I_NaR,_ in contrast, is revealed on membrane *hyperpolarizations* from the depolarized membrane potentials that evoke I_NaT_ (**Figure 1A**). In addition, the time courses of I_NaR_ activation and decay are much slower than I_NaT_ activation and decay (**Figure 1A**). Additional experiments revealed that, in response to membrane hyperpolarizations following brief (5 ms) depolarizing steps (to 0 mV), the amplitude of I_NaR_ varies as a function of the membrane potential of the hyperpolarizing voltage step (**Figure 1B**). The maximal amplitude of I_NaR_ is observed at approximately -45 mV (**Figure 1C**).

**Figure 1.**
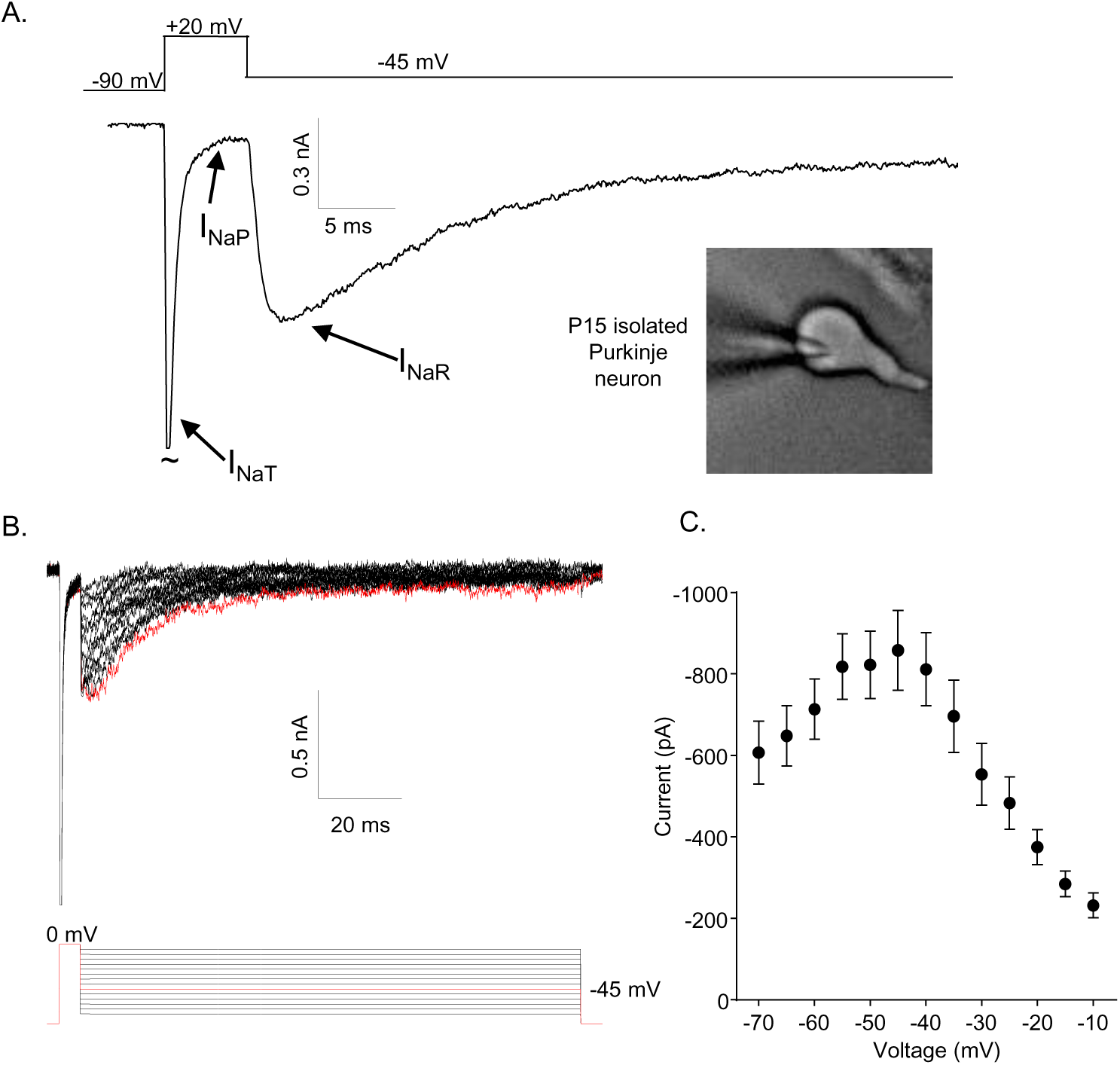
Mouse cerebellar Purkinje neurons express three Nav current components. (A) Representative recording of the transient (I_NaT_), persistent (I_NaP_), and resurgent (I_NaR_) components of the Nav currents in an isolated neonatal mouse cerebellar Purkinje neuron. The voltage-clamp paradigm is displayed above the current record, and the labelled arrows indicate the three Nav current components. (B) I_NaR_ waveforms, recorded during hyperpolarizing voltage steps to various potentials ranging from -70 mV to -10 mV, following 5 ms depolarizing voltage steps to 0 mV from a holding potential (HP) of -80 mV; the voltage-clamp paradigm is shown below the current records. The current record highlighted in red was recorded during the -45 mV hyperpolarizing voltage step (also indicated in *red* in the illustrated voltage-clamp paradigm). (C) Mean ± SEM (n = 15) peak I_NaR_ amplitudes are plotted as a function of the hyperpolarizing test potential; the peak I_NaR_ is recorded at approximately -45 mV.

To determine how the duration and the voltage of the depolarizing voltage step (that evokes I_NaT_) affect the amplitudes and waveforms of I_NaR_, voltage-clamp paradigms were designed in which either the duration or the voltage of the depolarizing step was varied (**Figure 2**). In initial experiments, the external and internal Na^+^ concentrations were 154 mM and 8 mM, respectively, resulting in a Na^+^ reversal potential of +75 mV. These experiments revealed that prolonging the duration of the +20 mV depolarizing voltage step resulted in the marked attenuation of the peak amplitudes of I_NaR_ measured during hyperpolarizing voltage steps to -45 mV (**Figure 2A**). The time course of the attenuation of I_NaR_ was well described by a single exponential characterized by a mean ± SEM (n = 12) time constant of 15.5 ± .5 ms (**Figure 2B**).

**Figure 2.**
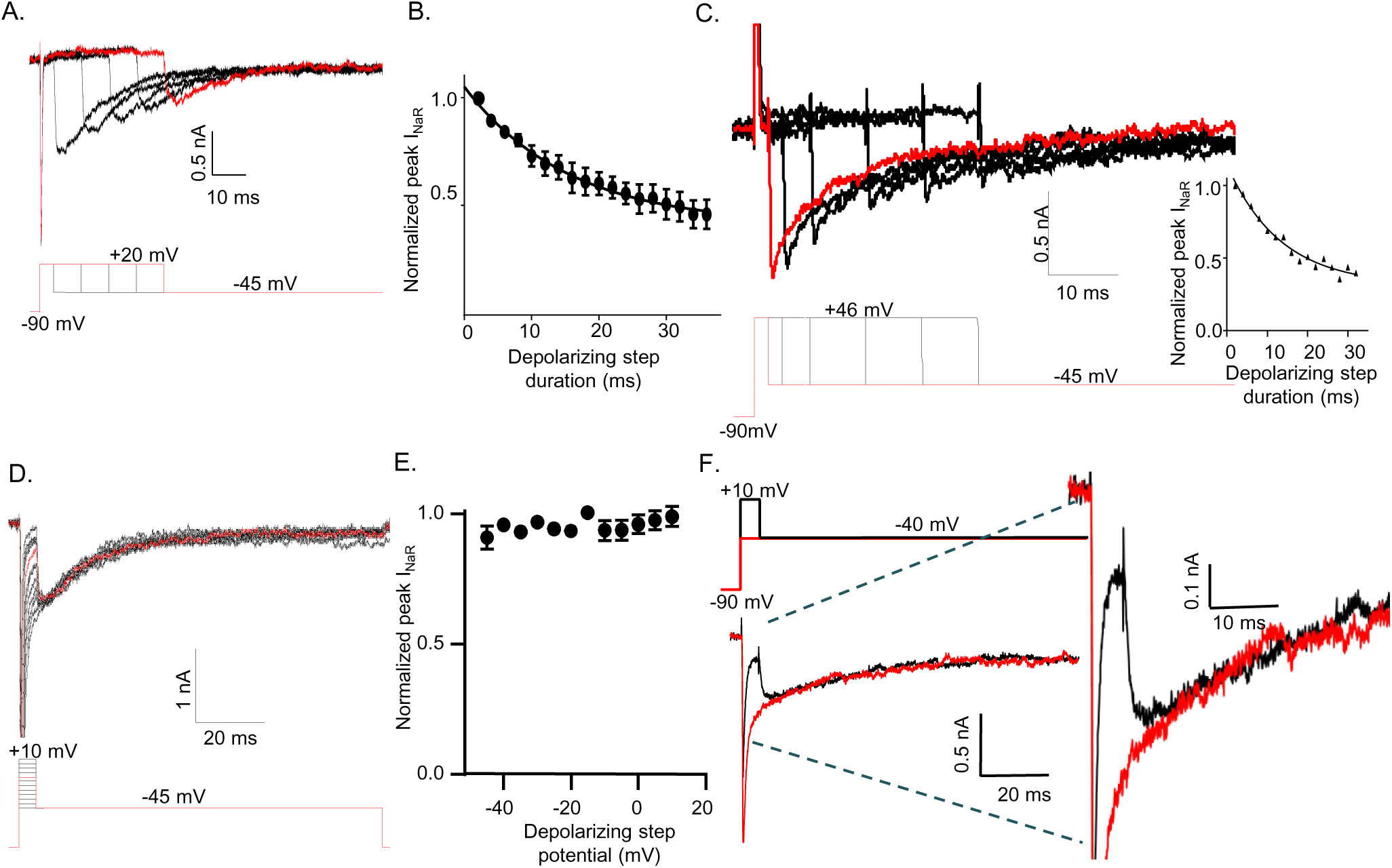
The amplitude of I_NaR_ is determined by the duration of the prior membrane depolarization. (A) In a neonatal mouse cerebellar Purkinje neuron, I_NaR_ was revealed on membrane hyperpolarizations following depolarizing voltage steps to +20 mV of varying durations; the voltage-clamp paradigm is shown below the current records. (B) Peak I_NaR_ amplitudes, evoked at -45 mV following each depolarizing voltage step to +20 mV, were measured and normalized to the maximal peak I_NaR_ (measured in the same cell). The mean ± SEM (n = 12) normalized peak I_NaR_ amplitudes are plotted as a function of the duration of the depolarizing voltage step. The attenuation of peak I_NaR_ as a function of the duration of the depolarizing voltage step was well described by a single exponential with a mean ± SEM time constant of 15.5 ± 0.5 ms (n = 12). (C) The dependence of I_NaR_ on the duration of the depolarizing voltage step was also measured with reduced (50 mM) extracellular and increased intracellular (15 mM) sodium, resulting in a Na^+^ reversal potential of +30 mV. Under these conditions, depolarizing voltage steps to +46 mV evoked outward I_NaT_. Peak I_NaR_ amplitudes, revealed during hyperpolarizing voltage steps to -45 mV, however, were also found to vary as a function of the duration of the depolarizing voltage step, revealing that I_NaR_ and the time-dependent attenuation of I_NaR_ are not affected by the direction (inward versus outward) of Na^+^ flux during the depolarizing voltage step. The peak amplitudes of I_NaR_, evoked at -45 mV following each depolarizing voltage step, were measured in each cell and normalized to the maximal I_NaR_ amplitude (in the same cell). As is evident from the representative records and the plot of normalized peak I_NaR_ amplitudes (on the right), the attenuation of I_NaR_ as a function of the duration of the depolarizing voltage steps to +46 mV is also well-described by a single exponential with a mean ± SEM time constant of 13.8 ± 1.1 ms (n = 6), a value similar to that observed when I_NaT_ was inward (B). (D) Representative I_NaR_ waveforms, recorded directly on repolarizations to -45 mv following 5 ms depolarizations to various membrane potentials from a -90 mV HP, are shown; the voltage-clamp protocol is shown below the current records. (E) The mean ± SEM (n = 6) normalized peak I_NaR_ amplitudes are plotted as a function of potential of the depolarizing voltage step. (F) Representative voltage-clamp recordings of Nav currents evoked (in the same cell) on direct depolarization to - 40 mV from a HP of -90 mV (*red*) and on hyperpolarization to -40 mV following a 5 ms depolarizing voltage step to +10 mV *(black*) from the same HP; the voltage-clamp protocols are shown above the current records and the currents are shown on an expanded scale on the right.

Subsequent experiments explored the effect of the driving force on Na^+^ on the time-dependent attenuation of I_NaR_. In these experiments, the extracellular Na^+^ was reduced to 50 mM and the Na^+^ concentration in the internal solution was increased to 15 mM, resulting in a Na^+^ reversal potential of approximately +30 mV. Under these recording conditions, depolarizations to +46 mV resulted in outward I_NaT_ (**Figure 2C**). Similar to the results obtained with inward I_NaT_ (**Figure 2A,B**), prolonging the depolarizing (+46 mV) voltage step when I_NaT_ is outward results in the rapid attenuation of the amplitudes of I_NaR_ evoked during the subsequent hyperpolarizations to -45 mV (**Figure 2C**). Under these conditions (outward I_NaT_), the time course of the attenuation of I_NaR_ was also well described by a single exponential with a mean ± SEM (n = 6) time constant of 13.8 ± 1.1 ms (**Figure 2C**), a value that is very similar to that observed when I_NaT_ is inward (**Figure 2B**). Taken together, these combined results demonstrate that, following brief depolarizations, there is a time-dependent accumulation of Nav channels (which underlie I_NaR_) in a non-conducting state, and that this accumulation occurs independent of the direction of the movement of permeating Na^+^ ions.

To determine how the voltage of the depolarizing step affects the amplitudes and kinetics of I_NaR_, the currents recorded at -45 mV after 5 ms depolarizing steps to various membrane potentials (**Figure 2D**) were measured. These experiments revealed that hyperpolarizations to -45 mV following brief (5 ms) depolarizations to various membrane potentials (ranging from -45 mV to +10 mV) resulted in identical I_NaR_ amplitudes (**Figure 2E**). Additionally, varying the voltage of the depolarizing step did not affect the kinetics of the decay of I_NaR_ (**Figure 2D**). This is clearly illustrated in **Figure 2F**, in which Nav currents recorded (in the same cell) at -40 mV during a sustained voltage step (*red*) and following a 5 ms depolarizing voltage step to +10 mV are superimposed. The opening of Nav channels that conduct I_NaR_, therefore, is not affected by the voltage of the prior membrane depolarization. Taken together, these observations suggest that there are two parallel, and kinetically distinct, Nav channel inactivation pathways: a fast inactivating pathway that is responsible for I_NaT_; and, a second, slower inactivation pathway that undelies I_NaR_.

### A novel Markov model, with parallel inactivation pathways, for Nav channel gating in Purkinje neurons

The results of the voltage-clamp experiments described above suggest that there are (at least) two distinct inactivation pathways that contribute to the gating of the Nav channels expressed in mouse cerebellar Purkinje neurons, i.e., one that is populated quickly on channel opening and inactivates rapidly (fast inactivation), and a second that is populated and decays much more slowly (slow inactivation). To explore this hypothesis, we developed a Markov kinetic state model that, after numerical optimization (see: **Methods**), recapitulates the range of time- and voltage-dependent properties observed experimentally for the Nav currents in mouse cerebellar Purkinje neurons. The optimized Markov model (**Figure 3A**) includes parallel fast (IF1, IF2) and slow inactivation (IS) pathways to reconcile the experimental findings (**Figure 2**) that the duration of the depolarizing voltage step that underlie Nav channel activation, and not the potential of the depolarizing voltage step or the direction (i.e., inward or outward) of the Na^+^ flux through open channels, determines the amplitudes of I_NaR_ recorded during subsequent membrane hyperpolarizations. The model was constrained to fit the experimental data derived from multiple voltage-clamp protocols designed to detail the properties of I_NaT_, including those to determine the voltage-dependence of I_NaT_ activation and steady-state inactivation, the time course of I_NaT_ recovery from inactivation (**Figure 3B-D**), and the time constant (tau) of decay of the peak I_NaT_. The model was further constrained by the experimental data obtained using protocols designed to detail the properties of I_NaR_, including the voltage-dependence of the ratio of the amplitudes of I_NaR_ and I_NaT_ (I_NaR_:I_NaT_), the time-dependent attenuation of I_NaR_ amplitudes, observed as a function of the duration of the depolarizing voltage step (**Figure 2A** and **B**), and the time constants (tau) of I_NaR_ decay determined for the currents recorded during hyperpolarizing voltage steps to various membrane potentials (**Figure 3E-G)**. Consistent with the experimental results presented in **Figure 2**, simulations using this gating model reveal that the amplitude of I_NaR_ is dependent on the duration, but *not* on the potential, of the depolarizing voltage step that evokes I_NaT_ (**Figure 3F, H, I**).

**Figure 3.**
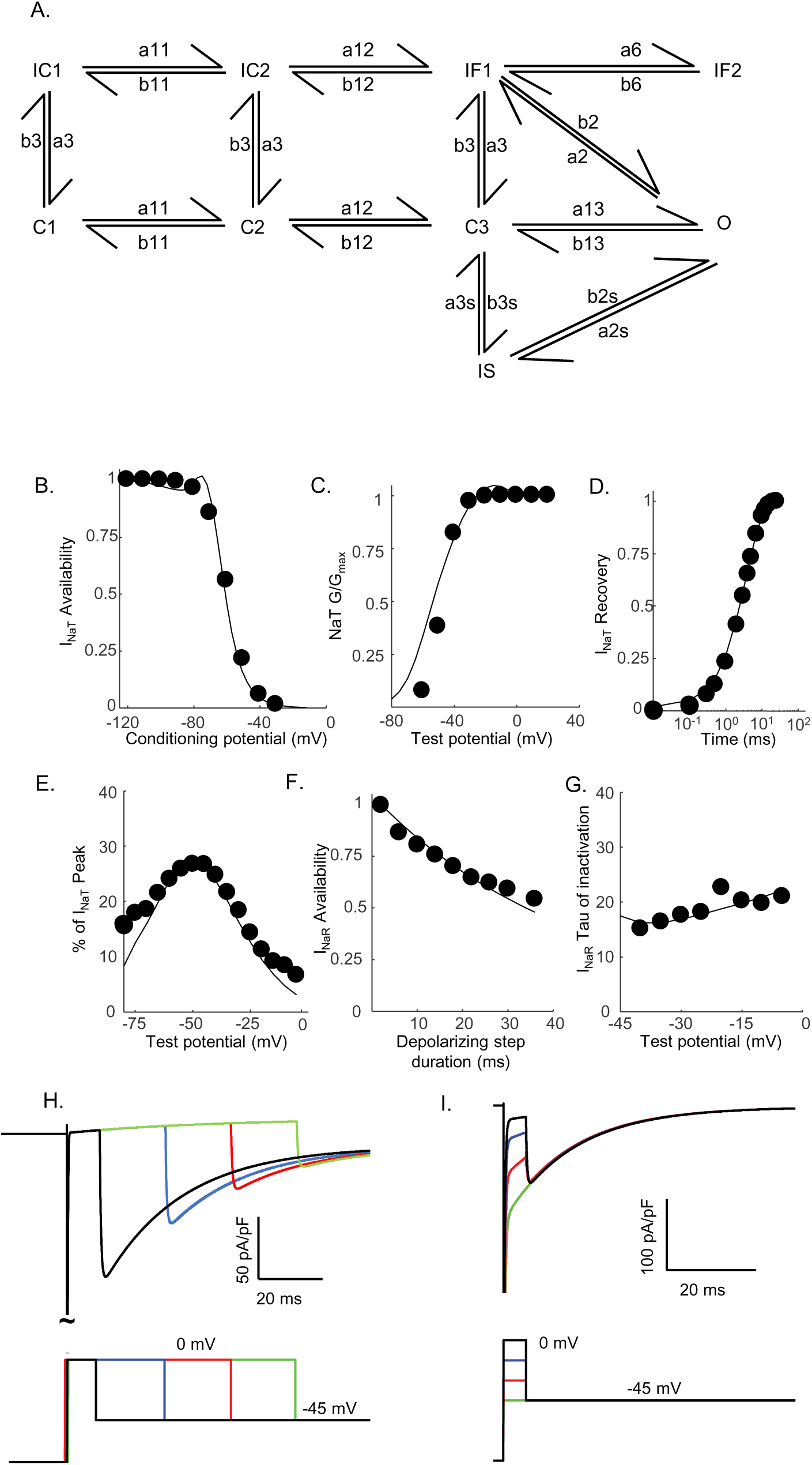
Novel Markov kinetic state model of Nav channel gating in cerebellar Purkinje neurons. A novel Markov kinetic state model was developed with parallel fast inactivating (IF1, IF2) and slow inactivating (IS) gating pathways (A). The model was numerically optimized (see: **Methods**) by simulating the data generated using voltage-clamp protocols identical to those used in the experiments to determine the detailed time- and voltage-dependent properties of the Nav currents in cerebellar Purkinje neurons. The various rate constants in the model were numerically optimized to recapitulate the measured properties of I_NaT_, I_NaP_, and I_NaR_ including the voltage-dependences of steady-state inactivation (B) and activation (C) of I_NaT_, and the kinetics of I_NaT_ recovery from inactivation (D). The model also reproduces the measured properties of I_NaR_, including the magnitude of I_NaR_ relative to I_NaT_ (E), the attenuation of the peak I_NaR_ amplitude as a function of the duration of the depolarizing voltage steps (F), and the kinetics of the decay (inactivation) of peak I_NaR_ amplitudes (G). Filled circles represent the mean experimental data and the lines represent the results of the simulation. The model successfully reproduces the observed, time-dependent attenuation of peak I_NaR_ amplitudes that is evident experimentally on membrane hyperpolarizations following depolarizing voltage steps of varying durations (H), and the finding that the peak amplitude of I_NaR_ is not affected by the potential of the depolarizing voltage step (I).

One potential benefit of computational modeling is the ability to dissect out possible mechanisms of channel gating by examining the occupancy of the individual channel states as a function of voltage and time. Taking advantage of this benefit, we examined the proportion of Nav channels populating each gating state during a simulated voltage-clamp protocol that evoked I_NaR_ at -45 mV after a 5 ms depolarizing voltage step to 0 mV (**Figure 4**). As illustrated, fast inactivation of I_NaT_ (during the 0 mV step) reflects (simulated) channels exiting the open state and accumulating into the IF1/IF2 states. The activation of I_NaR_ (at -45 mV) reflects channel transitioning from IF1/IF2 back into the open state, and the decay of I_NaR_ during the -45 mV step reflects the time-dependent accumulation of (simulated) Nav channels in the secondary, slow-inactivated state, IS (**Figure 3A**). The distinct pathways of inactivation are separated in time, but are not distinguished by differing voltage-dependences. Additionally, **Supplemental Figure 1** shows that simulated channels follow a similar inactivation pathway during a sustained depolarization to 0 mV, with an initial accumulation in the IF1/IF2 states and subsequent accumulation in the IS state, a property of the model that reveals why I_NaR_ amplitudes are reduced as the duration of the depolarizing voltage step (that evokes I_NaT_) is increased (**Figure 3F, H**).

**Figure 4.**
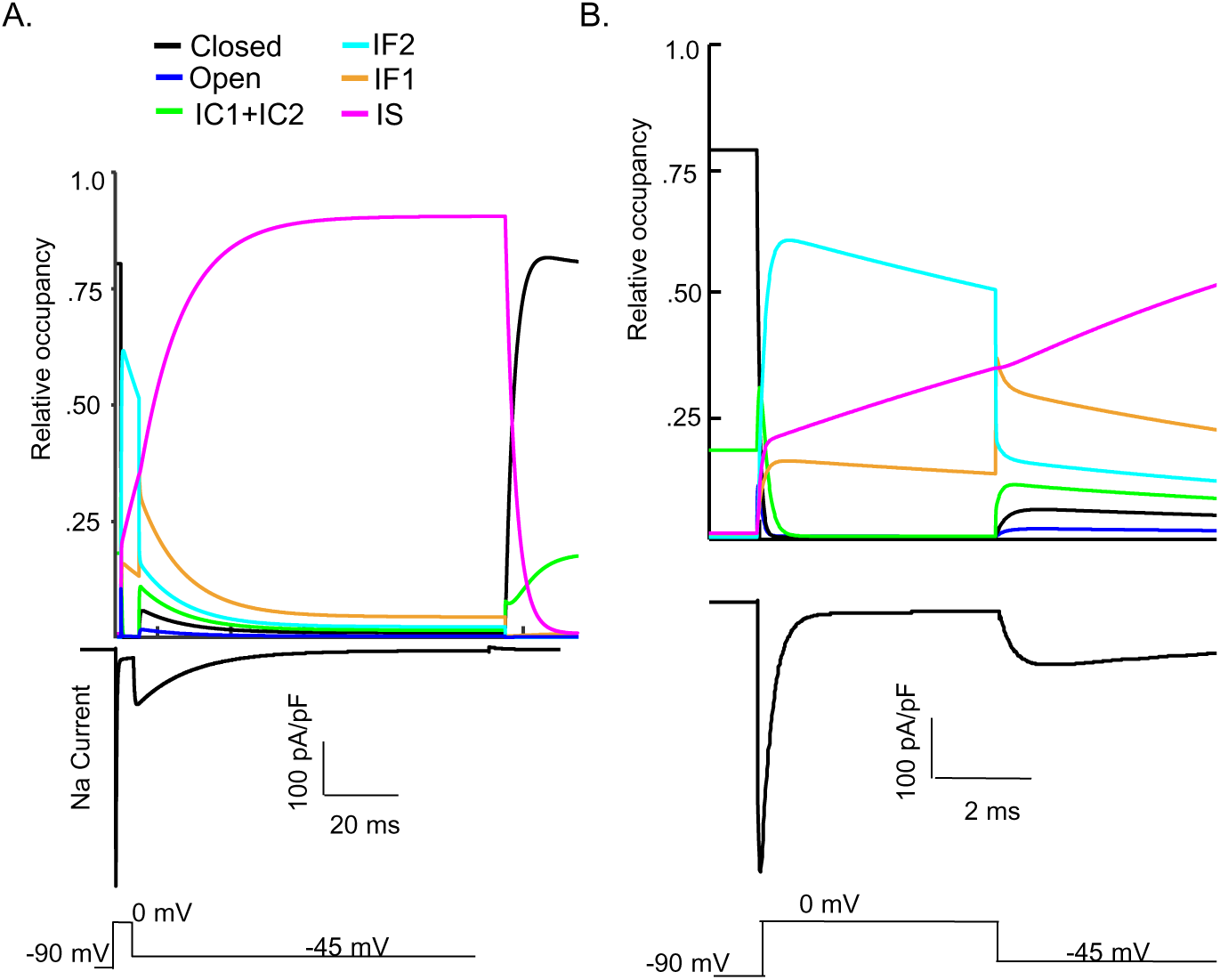
Kinetic state transitions during voltage-clamp simulations that evoke I_NaR_. There are two parallel pathways of Nav channel inactivation (IF1/IF2 and IS) in the novel Markov kinetic state model developed here (Figure 3). The occupancies of these states and of the other (i.e., closed, open etc.) channel gating states during a simulated voltage-clamp protocol, in which I_NaR_ is revealed on membrane hyperpolarization to -45 mV following a brief (5 ms) depolarizing voltage step to 0 mV from a holding potential of -90 mV, are plotted as a function of time in (A). The simulated voltage-clamp records and the experimental paradigm are illustrated below the gating state occupancy plot. Expanded (in time) views of the gating state occupancies and the simulated Nav currents are presented in (B).

### I_NaT_ and I_NaR_ are differentially sensitive to entry into the slow-inactivated state

The simulations (**Figure 4**) suggest that the fast decay of I_NaT_ and the much slower decay of I_NaR_ reflect separate, i.e., fast and slow, pathways of Nav channel inactivation and, in addition, that the decay of I_NaR_ reflects Nav channels accumulating in an absorbing, slow-inactivated state (i.e., the IS state in **Figure 3**). To test this hypothesis directly, we measured peak I_NaT_ and I_NaR_ amplitudes, recorded at 0 mV and -45 mV, respectively, during sequential voltage-clamp protocols separated by a brief (20 ms) interval at -90 mV; the paradigm is illustrated in **Figure 5A** below the current records. The 20 ms interval at -90 mV between the sequential protocols was determined to be sufficient for the near complete recovery of I_NaT_ from fast inactivation (Ransdell et al., 2017; Aman and Raman, 2007). However, if the channels that underlie I_NaR_ have accumulated in the second, slow-inactivated (IS) state during the first -45 mV voltage step and recovery from this state is also slow, one would expect to see differential effects on peak I_NaT_ and peak I_NaR_ amplitudes when the time interval at -90 mV is sufficient to allow complete recovery of I_NaT_, but too short to allow the complete recovery of I_NaR_. As illustrated in the representative records shown in **Figure 5A**, this voltage-clamp paradigm revealed that, when the time interval at -90 mV was reduced to 20 ms, the amplitude of I_NaR_ was indeed reduced to a greater extent than the amplitude of I_NaT_. Recordings from five additional Purkinje neurons using this voltage-clamp paradigm yielded similar results. Plotting the relative peak amplitudes of I_NaT_ and I_NaR_ measured during the second protocol, compared with the first, reveals that the 20 ms hyperpolarizing voltage step to -90 mV was sufficient to provide nearly complete (0.95 ± .01; n = 6) recovery of I_NaT_ (**Figure 5B**), whereas there was a marked reduction in the amplitude of I_NaR_ (0.63 ± .05; n = 6), measured during the second, compared with the first, protocol (**Figure 5B**).

**Figure 5.**
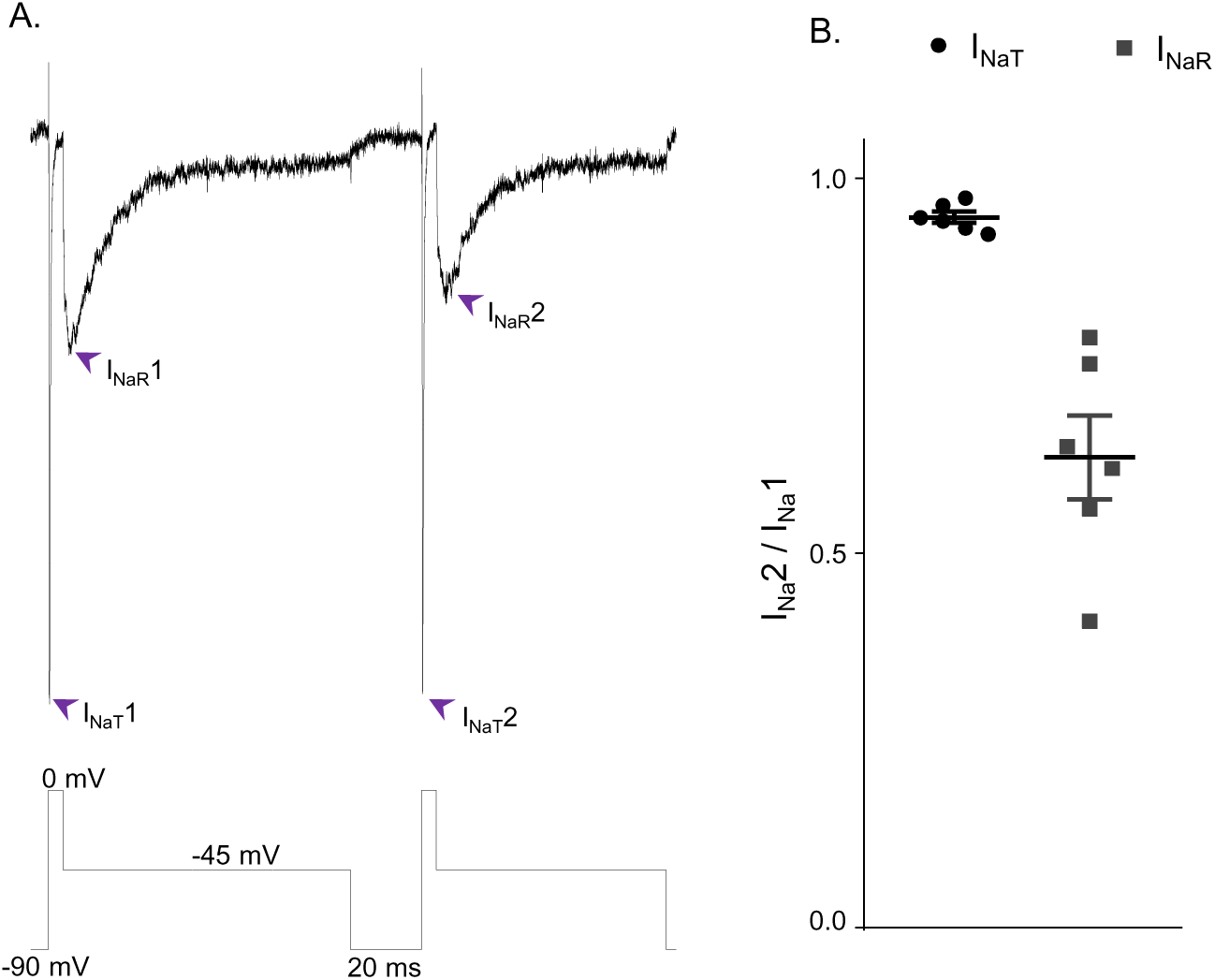
I_NaR_ and I_NaT_ display distinct rates of recovery from inactivation. (A) Representative Nav currents recorded in a mouse cerebellar Purkinje neuron during a voltage-clamp paradigm designed to determine if the relative rates of recovery from inactivation of I_NaT_ and I_NaR_ are distinct. Inward Nav currents were recorded during sequential and identical voltage-clamp steps (to 0 mV for 5 ms and to -45 mV for 100 ms), separated by a brief (20 ms) hyperpolarizing step to -90 mV; the voltage-clamp paradigm is shown below the current records. As is evident, the amplitude of I_NaR_ (at -45 mV) during the second voltage-clamp step to -45 mV was attenuated more than I_NaT_ (during the second step to 0 mV). Similar results were obtained in five additional Purkinje neurons using the voltage-clamp paradigm shown. (B) Plot of the relative peak I_NaT_ (circles) and peak I_NaR_ (squares) amplitudes measured during the second voltage-clamp steps (to 0 mV and -45 mV), compared with the first. As is evident, the relative amplitude of I_NaR_ is reduced (0.63 ± .05; n = 6) to a greater extent (paired Student’s t-test; *P* ≤ .01) than I_NaT_ (0.95 ± .01; n = 6). The mean ± SEM (n = 6) relative I_NaT_ and I_NaR_ amplitudes are also indicated.

### I_NaR_ reflects the transitioning of fast-inactivated Nav channels into an open conducting state

The results presented in **Figure 5** indicate that Nav channels open during membrane hyperpolarizations from depolarized potentials and that these (open) channels undergo inactivation at a rate that is clearly distinct from the rapid decay of I_NaT_. To determine the relationship between I_NaR_ and the channels that give rise to the persistent component of the sodium current, I_NaP_, we used two voltage-clamp protocols, designed to allow direct measurements of I_NaP_ alone or I_NaP_ plus I_NaR_. In the first protocol, a slow (dV/dt = 0.12 mV/ms) depolarizing voltage ramp (from -100 mV to 0 mV) was presented and inward currents, reflecting only I_NaP_, were recorded (**Figure 6A**, *blue*). In the same cell, we also recorded Nav currents evoked during a slow (dV/dt = 0.12 mV/ms) hyperpolarizing (from 0 mV to -100 mV) voltage ramp (**Figure 6A**, *red*). In the latter case, the measured inward currents reflect both I_NaP_ and I_NaR_. In addition, we recorded the currents evoked during depolarizing voltage steps to various test potentials between -75 mV and +10 mV from a holding potential of -100 mV, and we measured the amplitudes of I_NaP_ directly, at 25 ms after the onset of each depolarizing voltage step (**Figure 6A**, green). The current-voltage plots, derived from the data obtained in these experiments, are presented in **Figure 6B**; the colors correspond to those used to illustrate the current records presented in **Figure 6A**. As is evident, the voltage-dependences and the magnitudes of the Nav currents recorded using the three voltage-clamp protocols (in the same cell) are indeed indistinguishable. Similar results were obtained in recordings from 4 additional Purkinje neurons (see: **Discussion**).

**Figure 6.**
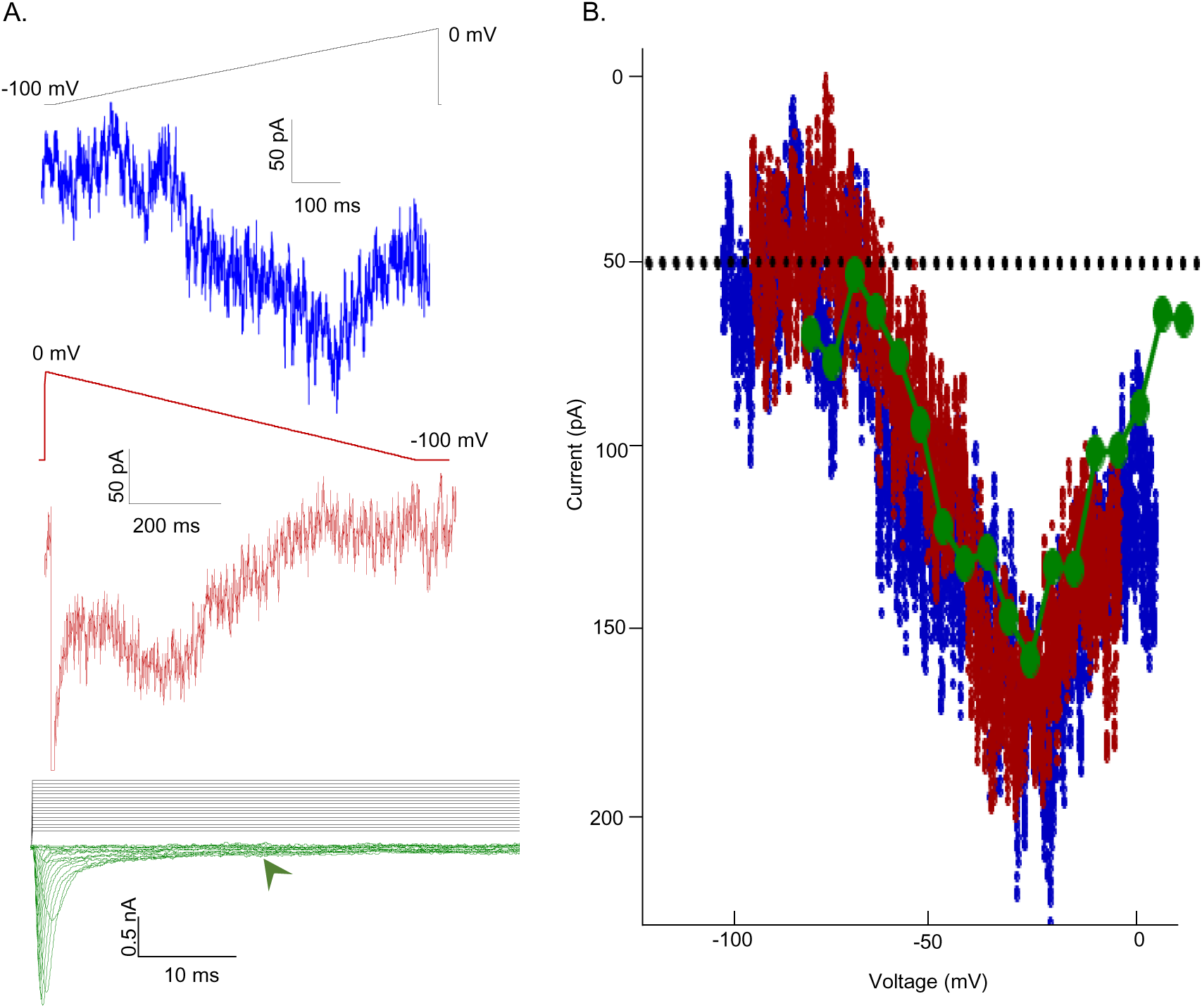
Voltage-dependences of activation of I_NaR_ and I_NaP_ are indistinguishable. (A) To test the hypothesis that non-inactivating Nav channels underlie I_NaR_, a depolarizing voltage ramp (*blue*) protocol (from -100 mV to 0 mV at 1.2 mV/ms) and a steady-state voltage step (*green*) protocol (with depolarizations from a holding potential of -100 mV to test potentials ranging from -75 to 10 mV in 5 mV increments) were used to reveal the magnitude and voltage-dependent properties of the non-inactivating (persistent) component of the Nav current, I_NaP_. In addition, a hyperpolarizing voltage ramp (from 0 mV to -100 mV at 1.2 mV/ms or dV/dt), presented after a 5 ms depolarizing step to 0 mV, was used to reveal both I_NaR_ and I_NaP_. Note that the three representative records shown were obtained from the same Purkinje neuron. (B) The current-voltage relationships, derived from the records presented in (A) are plotted (in the corresponding color). From the records shown in the lowest panel of (A), the amplitudes of the steady state inward currents at 25 ms at each test potential are plotted as points (green); the current amplitudes determined (at 2 ms intervals) from the ramp protocols (*red* and *blue* traces) are also plotted. As is evident, the current-voltage relations of the Nav currents recorded using the three different voltage-clamp protocols overlap; the magnitudes of the inward Nav currents are also indistinguishable. Similar results were obtained in 4 additional Purkinje neurons.

### Nav channel gating model with an open-blocked state does not reproduce the voltage-clamp data

The novel Markov model developed here (**Figure 3A**) is quite different from the previously proposed model of Nav channel gating in mouse cerebellar Purkinje neurons (Raman and Bean, 2001). In this earlier model (illustrated in **Figure 7A**), there are two distinct competing pathways that depopulate the open state, one of which involves fast Nav channel inactivation and results in the population of the I6 state (**Figure 7A**) and the other, competing, pathway involves the blockade of open Nav channels and the generation of the open-blocked (OB) state (**Figure 7A**). The isolation of the open-blocked state from all of the other kinetic states except the open state is a distinctive feature in the model of Raman and Bean (**Figure 7A**). This configuration means that channels that have entered the OB state can only exit this state (i.e., become unblocked) by transitioning into the open/conducting state to generate resurgent Na^+^ influx, i.e., I_NaR_ (Raman and Bean, 2001). Although it was suggested that the blocking particle responsible for producing the OB state was a protein, specifically the Nav channel accessory subunit Navβ4 (Grieco et al., 2002; Grieco et al., 2005; (Grieco et al., 2005, Bant and Raman, 2010), it was subsequently demonstrated that I_NaR_ is reduced, but is *not* eliminated, in cerebellar Purkinje neurons in (*Scn4b^-/-^*) mice lacking Navβ4 (Ransdell et al., 2017; White et al. (2019) (see: **Discussion**).

**Figure 7.**
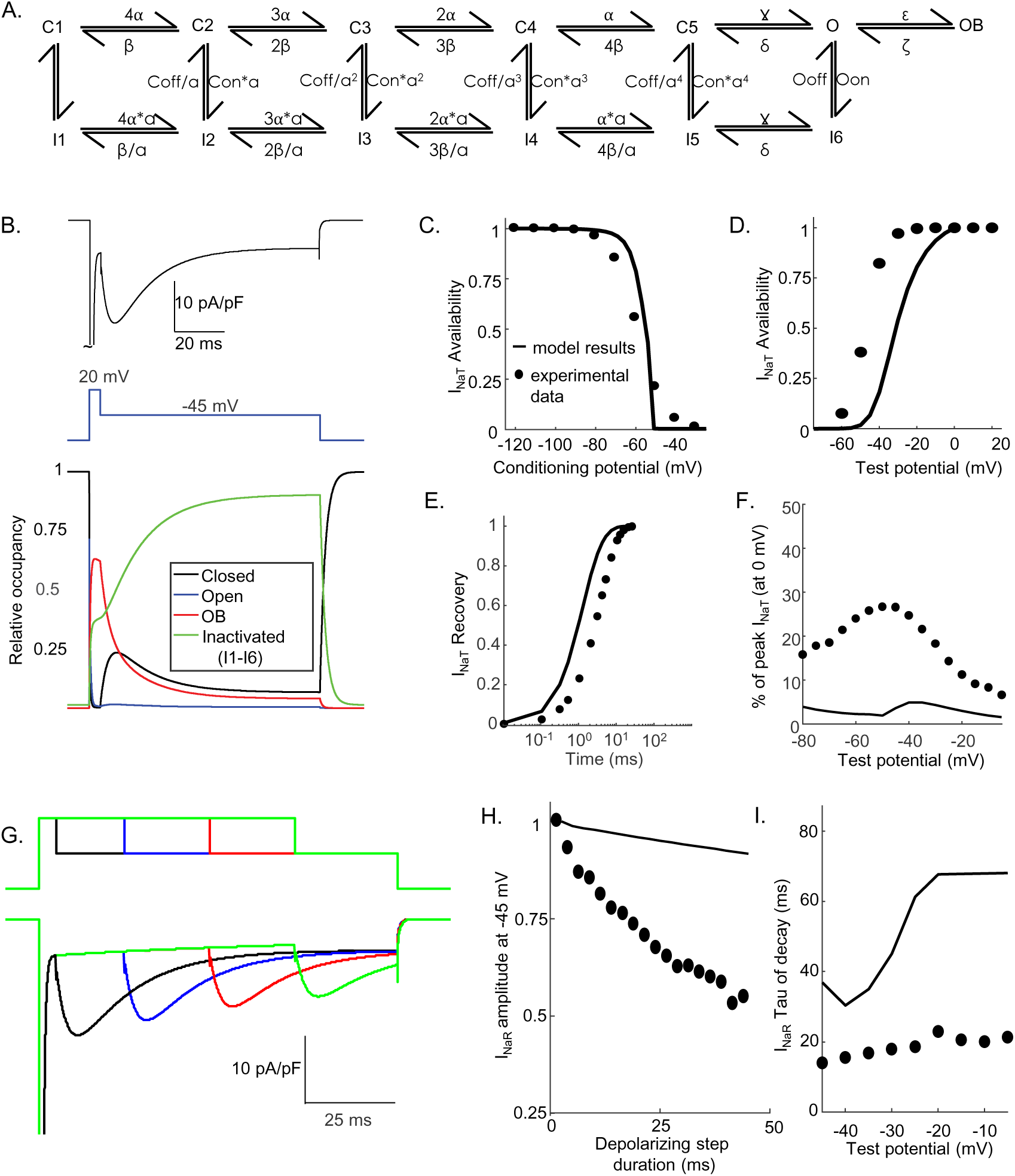
Simulations using the Raman-Bean model of Nav channel gating do not recapitulate the acquired voltage-clamp data. (A) The previously described Markov kinetic state model of Nav channel gating in mouse cerebellar Purkinje neurons (Raman and Bean, 2001) is illustrated. (B) Representative simulated inward Nav current waveforms, produced by this (A) model using the voltage-clamp paradigm shown below the current records, are presented. The time-matched normalized occupancies of the combined closed (C1-C5), open (O), open-blocked (OB) and combined inactivated (I1-I6) gating states are plotted below the voltage protocol. (C-E) Comparisons of the time- and voltage-dependent properties of I_NaT_ derived from simulations using the model in (A) with (our) experimental data obtained in recordings from mouse cerebellar Purkinje neurons (the same data as were used to generate the results in Figure 3); filled circles represent the mean experimental data and the lines represent the results of the simulations. (F) Relative I_NaR_ amplitudes (normalized to peak I_NaT_ at 0 mV) are plotted as a function of the hyperpolarizing test potential. (G) Simulated I_NaR_ waveforms, produced on membrane hyperpolarizations to -45 mV following depolarizing voltage steps to +20 mV of varying durations, are shown. (H) Peak normalized I_NaR_ amplitudes (at -45 mV), derived from the simulations in (G), are plotted as a function of the duration of the prior +20 mV depolarizing voltage step (solid line), together with the mean experimental data (filled circles) obtained in recordings from mouse cerebellar Purkinje neurons (the same data as used to generate the results in Figure 3). (I) The kinetics of I_NaR_ decay, derived from single exponential fits to the decay phases of the currents recorded at various membrane potentials, are presented in (I); the solid lines indicate the results of the simulations, and the filled circles are the mean experimental data obtained in recordings from mouse cerebellar Purkinje neurons (the same data as used to generate the results in Figure 3).

Subsequent efforts here were focused on determining directly whether the Nav channel gating model in which I_NaR_ is generated by the open-blocked mechanism (**Figure 7A**) could/would also reliably reproduce the detailed time- and voltage-dependent properties of the Nav currents determined experimentally in mouse cerebellar Purkinje neurons (and presented in **Figures 1, 2, 5** and **6**). As illustrated in **Figure 7B**, simulations with the open-blocked model for I_NaR_ generation (**Figure 7A**) provided transient and resurgent Nav current components (**Figure 7B, upper panel**) that resemble those measured experimentally (**Figure 1A**). A time-locked plot of gating state occupancies with this model reveals that I_NaR_ activation occurs as simulated channels exit the OB state and into the open (conducting) state (**Figure 7B, lower panel**). Additional simulations revealed that this model also recapitulates the voltage- and time-dependences of I_NaT_ activation, inactivation and recovery from inactivation (**Figure 7C-E**) in mouse cerebellar Purkinje neurons, although the modelled I_NaT_ activates at more hyperpolarized voltages than native I_NaT_. Although recapitulating the voltage-dependence of I_NaR_ activation (**Figure 7F**), the model also predicts that the magnitude of I_NaR_ evoked at all hyperpolarized membrane potentials, relative to the peak amplitude of I_NaT_ (evoked at 0 mV), is much smaller than observed experimentally (**Figure 7F**).

In the open-channel block model of I_NaR_ gating (**Figure 7A**), more positive depolarizing voltage steps promote entry into the open-blocked state, whereas channels are favored to undergo fast (conventional) inactivation at more hyperpolarized membrane potentials (Raman and Bean, 2001, Lewis and Raman, 2014). When the voltage-clamp protocols used to generate the data presented in **Figure 2** were used in simulations with the open-channel block model (**Figure 7A**), however, substantial differences between the predictions of this model and our experimental data were revealed. In contrast to what is observed experimentally (**Figure 2**), for example, increasing the duration of the brief (5 ms) depolarizing voltage step (to 0 mV) in the open channel block model resulted in very little time-dependent attenuation of the amplitude of I_NaR_ recorded on membrane hyperpolarization to -45 mV (**Figure 7G, H**). In addition, the time course of the decay of the resurgent currents predicted by the open channel block model (**Figure 7A**) are much slower than we observed experimentally for I_NaR_ in mouse cerebellar Purkinje neurons (**Figure 7I**).

### The fast and slow-inactivated Nav channel states are populated separately and at different rates

The experimental data and the simulations using the novel Nav channel gating model developed here (**Figure 3A**) suggest that I_NaR_ is mediated by Nav channels transiting from a fast-inactivated state (IF1) into the open state, and subsequently accumulating in an absorbing, slow-inactivated state (IS). It is also possible, however, that the absorbing slow-inactivated state that underlies I_NaR_ decay reflects Nav channels accumulating into a long-term inactivated state, i.e., a state in which channels are non-conducting for hundreds of ms, for example by a blocking particle(s) that competes on a time-scale similar to conventional fast inactivation (Dover et al., 2010, Venkatesan et al., 2014). To test this possibility directly, a voltage-clamp protocol was developed to allow direct comparison in the same cell) of I_NaR_ recorded during a single (80 ms) hyperpolarizing voltage step to -45 mV, presented following a brief (5 ms) depolarization to 0 mV, with I_NaR_ recorded at -45 mV during successive brief (2 ms) hyperpolarizing voltage steps interspersed with brief (5 ms) depolarizations to 0 mV (**Figure 8A**). If a competing extrinsic blocking particle has fast-onset, competes with conventional inactivation and is absorbing, one would expect to see reductions in the amplitudes of I_NaR_ recorded during each successive hyperpolarizing voltage step to -45 mV compared with I_NaR_ recorded during a sustained hyperpolarizing voltage step to -45 mV. As is evident in the experimental records shown in **Figure 8A**, however, I_NaR_ waveforms evoked (in the same cell) using these two voltage-clamp protocols were quite similar.

**Figure 8.**
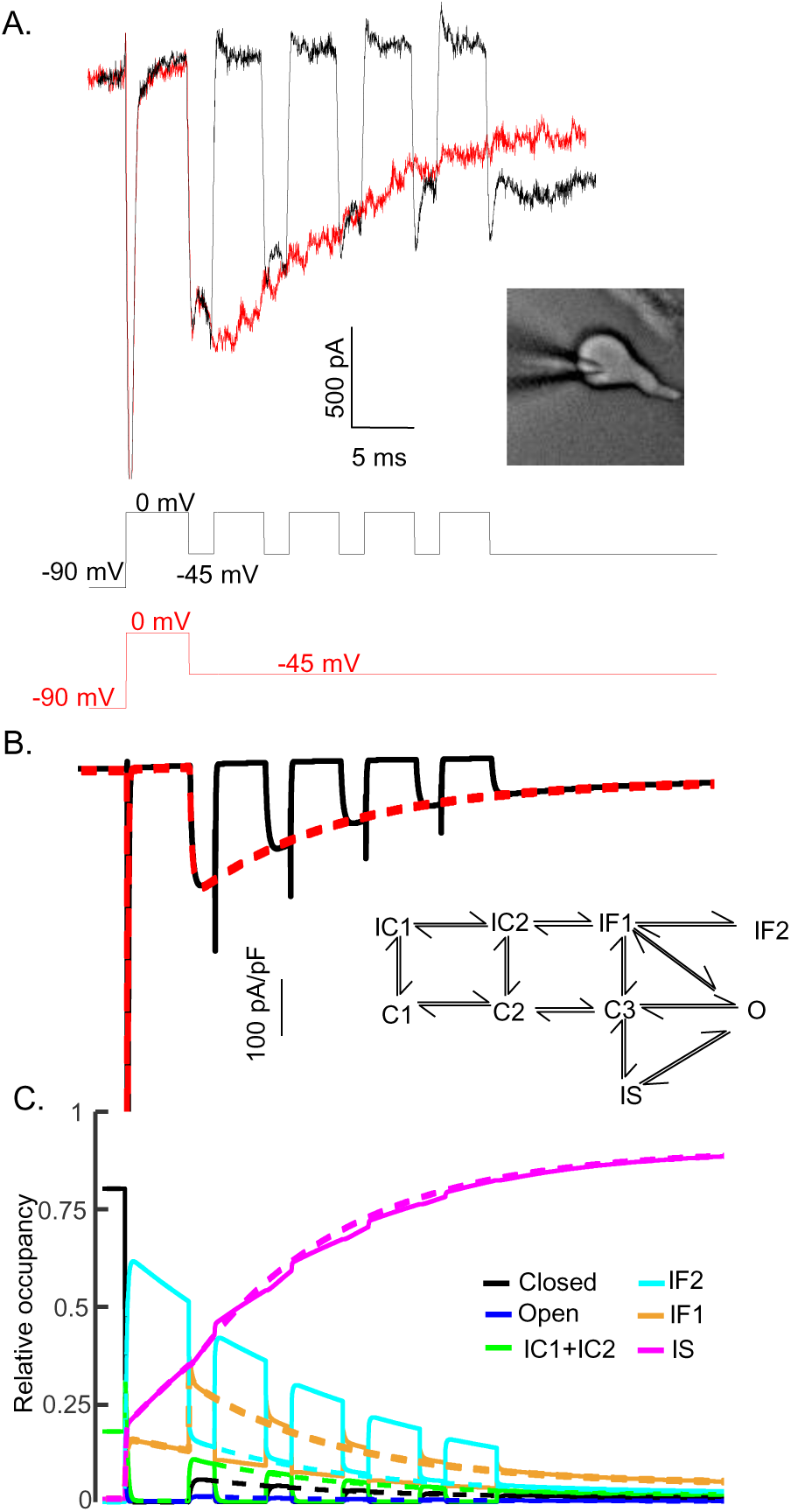
The time course and amplitude of I_NaR_ are recapitulated during repetitive brief depolarizing steps. To determine if two competing inactivation states underlie the observed differences in I_NaT_ and I_NaR_ recovery from inactivation (illustrated in Figure 5), a protocol was developed to allow direct comparison of I_NaR_ recorded during a single (80 ms) hyperpolarizing voltage step to -45 mV (*red*), presented following a brief (5 ms) depolarization to 0 mV, with I_NaR_ recorded (in the same cell) at -45 mV during successive brief (2 ms) hyperpolarizing voltage steps interspersed with brief (5 ms) depolarizations to 0 mV (*black*). Representative records are shown in (A); the voltage-clamp paradigms are illustrated below the current records. As is evident (A), the envelope of the currents generated using these two protocols superimpose, suggesting that the inactivation pathway responsible for I_NaR_ decay does not compete with fast inactivation. (B) Simulated current waveforms, generated using the same two voltage-clamp protocols illustrated in (A) with the novel kinetic-state model presented in Figure 3A, are shown. (C) Gating state occupancies for simulated current traces are shown. For direct comparison of the results of the simulations using the voltage-clamp protocols illustrated in (A) with the Raman Bean gating model (2001), see: **Supplemental Figure 2**.

The experimental data in **Figure 8A** were recapitulated in simulations (**Figure 8B**) using the novel gating state model (**Figure 3A**) developed here. The current waveforms generated by the model are indistinguishable from the experimental results (compare **Figures 8A** and **8B**). In addition, and without tuning any of the model parameters, the kinetic state occupancy plots generated using the two voltage-clamp protocols were also very similar (**Figure 8C**). This voltage-clamp protocol was also applied in simulations using the previously described (**Figure 7A**) open channel block model (Raman and Bean, 2001). In this case, in marked contrast with the results presented in **Figure 8**, the model does not reproduce the experimental data (**Supplemental Figure 2A**). The simulations revealed that, in this model, channels did not maximally enter into the OB state on depolarizations to 0 mV and the 2 ms hyperpolarizations were not sufficient to fully activate I_NaR_ (**Supplemental Figure 2B**). Additionally, in this model, during each of the successive depolarizations (to 0mV), transient Nav currents were revealed (**Supplemental Figure 2A**), reflecting channels exiting the OB state on membrane hyperpolarization and re-entering the OB state on membrane depolarization.

### Simulating I_NaR_ in Scn4b^-/-^ cerebellar Purkinje neurons

We previously reported that the targeted deletion of *Scn4b* in mice results in a marked (∼50%) reduction in I_NaR_ amplitudes in cerebellar Purkinje neurons (**Figure 9A**) without measurable effects on I_NaR_ waveforms (Ransdell et al., 2017). Similar results were subsequently reported in studies using a different *Scn4b^-/-^* mouse line (White et al., 2019). To explore the ability of the novel model of Nav gating, developed and presented here (**Figure 3A**), to scale the amplitude of I_NaR_ while leaving the time- and voltage-dependent properties of the currents unaffected, we optimized the parameters of the model (see: **Methods**) to reproduce the experimentally determined reduction in I_NaR_ amplitudes with the loss of Navβ4 (Ransdell et al., 2017; White et al., 2019). As illustrated in **Figure 9B**, although reduced in amplitude, the voltage dependence of I_NaR_ generated by the *Scn4b^-/-^* Nav channel gating model is very similar to wild type I_NaR._. In **Figure 9C**, representative I_NaR_ waveforms generated by the gating models of wild type and *Scn4b^-/-^* Nav currents are superimposed; the time courses of wild type and *Scn4b^-/-^* I_NaR_ are indistinguishable. Time-locked with the wild type and *Scn4b^-/-^* Nav current traces are plots of the Nav channel gating state occupancies as a function of time (**Figure 9D**) in the wild type (solid lines) and the *Scn4b^-/-^* (dashed lines) I_NaR_ models. The gating state occupancy plots (**Figure 9D**) reveal that the attenuation of the amplitude of I_NaR_ in the *Scn4b^-/-^* model is the result of the reduced accumulation of channels in the IF2 state during the initial depolarization and increased occupancy in the IS state. Together, these data suggest that Navβ4 delays entry into the slow-inactivated state (IS), allowing for greater recovery from conventional inactivation and thus, larger I_NaR_ amplitudes (see: **Discussion**).

**Figure 9.**
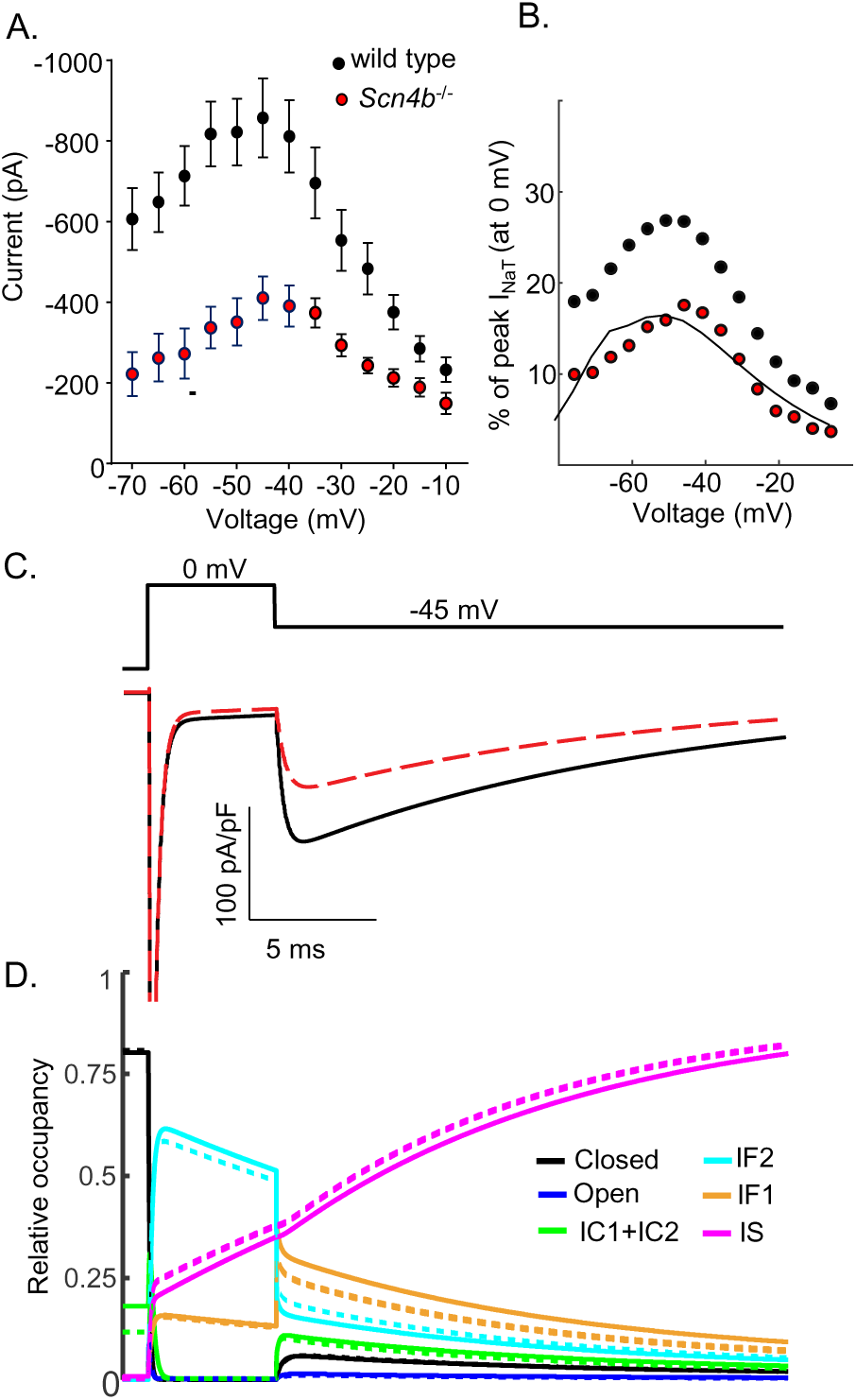
Promoting entry into the slow-inactivated state reduces I_NaR_ amplitudes. (A) Mean ± SEM peak I_NaR_ amplitudes, measured on membrane hyperpolarizations following brief depolarizing voltage steps to +10 mV, in wild type (*black*) and *Scn4b^-/-^* (*red*) mouse cerebellar Purkinje neurons are plotted as a function of membrane voltage are shown (data were reproduced with permission from: Ransdell et al., 2017). Peak I_NaR_ amplitudes in individual wild type and *Scn4b^-/-^* cells were also normalized to peak I_NaT_ measured (at 0 mV) in the same cell, and the mean I_NaR_ as a percentage of peak I_NaT_ in wild type (*black*) and *Scn4b^-/-^* (*red*) cells are plotted (as points) in (B); the solid line is the normalized relative I_NaR_/I_NaT_ generated by the *Scn4b^-/-^* model. (C) Consistent with the experimental data, the kinetics of I_NaR_ are not affected measurably by the loss of *Scn4b* (Navβ4) in the model, whereas I_NaR_ amplitudes are reduced to ∼50% of wild type I_NaR_ levels (C). A time-locked plot of the gating state transitions (D) indicates that I_NaR_ amplitudes are reduced in the *Scn4b^-/-^* model (dashed lines) due to a decrease in IF2 occupancy and an increase in IS occupancy.

## DISCUSSION

Using a combined experimental and modeling approach, we describe here a novel mechanism for I_NaR_ gating that reflects two kinetically distinct (fast and slow) pathways of Nav channel inactivation. Importantly, the model reliably recapitulates the detailed time- and voltage-dependent properties of I_NaR_ determined experimentally in mouse cerebellar Purkinje neurons. The model, for example, accounts for the experimental finding that the peak amplitude of I_NaR_ (recorded on membrane hyperpolarizations following brief depolarizations that evoke I_NaT_) is sensitive to the duration of the preceding depolarizing voltage step, with longer depolarizations resulting in lower I_NaR_ amplitudes. Importantly, the dependence of I_NaR_ amplitudes on the duration of the depolarizing voltage step was also observed when the inward driving force of Na^+^ was reduced or eliminated. In addition, the model reproduces the experimental finding that I_NaR_ amplitudes are not dependent on the voltage of the initial depolarizing voltage step (that evokes I_NaT_). Finally, and also consistent with the experimental data, the voltage-dependence and the amplitude of I_NaR_ (revealed on membrane hyperpolarizations) mirror the voltage-dependence and the magnitude of the non-inactivating or persistent Nav current, I_NaP_, component. Interestingly, this last observation, i.e., that the voltage-dependence of I_NaP_ is identical to the voltage-dependence as I_NaR_, was previously reported (Kay et al., 1998).

The model developed here (**Figure 3A**) is distinct from a previously proposed model (**Figure 7A**) of Nav channel gating in mouse cerebellar Purkinje neurons (Raman and Bean, 2001) that involves two competing pathways out of the open state, i.e., channels transitioning into either an open-blocked state or an inactivated state. In this earlier model (**Figure 7A**) of I_NaR_ gating, a blocking particle occludes the Nav channel pore that is opened on membrane depolarization, functionally competing with conventional, fast inactivation, and, in addition, the open-blocked state is isolated off the open/conducting state (Raman and Bean, 2001). In this model, entry into the open-blocked state is promoted by membrane depolarizations to positive potentials and, on subsequent membrane hyperpolarization, the blocking particle is expelled by the Na^+^ driven through the unblocked Nav channel pore (Aman and Raman, 2010). In addition, the larger the driving force on Na^+^, the more rapidly the blocker is displaced (Aman and Raman, 2010). This process, while intrinsically voltage-independent, therefore, is tied to the driving force on Na^+^, consistent with the experimental observation that the peak amplitude of I_NaR_ is observed at relatively hyperpolarized (-45 to -30 mV) membrane potentials (Aman and Raman, 2010; Lewis and Raman, 2014). At more negative membrane potentials, closed state inactivation begins to dominate, and I_NaR_ is reduced (Raman and Bean, 2001; Lewis and Raman, 2014). The experimental results presented here, however, reveal that the peak amplitude of I_NaR_ is not affected by the voltage of the initial membrane depolarization (**Figure 2D**), but rather *is affected by the duration* of the prior depolarizing voltage step (**Figure 2A, B**). In addition, experimental results presented here demonstrate that the time-dependent accumulation of Nav channels into the slow-inactivated state occurs independent of the direction of the movement of permeating Na^+^ ions.

### Recovery from conventional Nav channel inactivation into an open/conducting state

The experiments and the simulations presented here suggest that I_NaR_ reflects the transitioning of Nav channels that have undergone conventional fast inactivation, into an open/conducting state on membrane hyperpolarizations. Consistent with the slow decay of I_NaR_, the model includes two parallel, but distinct, inactivation pathways: the fast (i.e., IF1 and IF2) inactivation pathway and the slow (i.e., IS) inactivation pathway, which satisfies the key experimental finding that the amplitude of I_NaR_ is insensitive to the voltage of the initial depolarization (**Figure 2**). Although it was previously reported that the magnitude of I_NaR_ does not depend on the voltage of the initial depolarization, this observation was interpreted as suggesting that all Nav channels initially undergo open channel block (Raman and Bean, 2001). The results of the experiments here involving prolonged depolarizations (**Figure 2A-C**), however, are not consistent with this suggestion. In these experiments, we found that there is still a time dependent attenuation of I_NaR_ amplitudes on membrane hyperpolarizations following depolarizations to positive membrane potentials, with little or no inward driving force on Na^+^. These findings are inconsistent with the open-channel block hypothesis, in which positive membrane potentials are thought to promote and stabilize open-channel block (Raman and Bean, 2001; Aman and Raman, 2010; Lewis and Raman, 2014).

One assumption usually made when considering the gating of Nav channels is that deactivation must occur prior to recovery from inactivation. Or, to state another way, Nav channels cannot exit the conventional inactivated state directly into an open or conducting state. This idea is based on voltage-clamp experiments conducted on CA1 hippocampal neurons (Kuo and Bean, 1994) and the squid giant axon (Armstrong (2006), which revealed that recovery from inactivation occurs with a delay and is voltage-dependent. Indeed, if recovery from inactivation requires all four of the VSDs of domains I-IV to move into the deactivated position, the model presented here is not possible because channels that have undergone conventional, fast inactivation would not be able to move directly into an open/conducting state without first transiting through an intermediate closed state. However, if deactivation of VSD IV is sufficient to release the cytosolic DIII–DIV linker peptide from the channel pore, while VSD I, II and III remain in the activated conformation, Nav channels could transit from an inactivated state directly into an open/conducting state. There are several reports suggesting that this is possible. Schiavon et al. (2012), for example, demonstrated that scorpion beta toxins induce I_NaR_ in heterologously (in HEK-293 cells) expressed Nav channels. These toxins were previously shown to cause a negative-shift in the voltage-dependence of DI-VSD, DII-VSD and DIII-VSD activation by trapping the DII-VSD in the activated position after depolarizing prepulses (Cestele et al., 1998, Schiavon et al., 2006). Thus, on repolarization, the channel recovers from inactivation, allowing resurgent Na^+^ influx.

Resurgent Nav currents have also been induced in heterologously (in HEK-293 cells) Nav1.5-encoded channels following application of a classical type-II pyrethroid, deltamethrin. Similar to our model for the generation of I_NaR_, the deltamethrin-modified Nav1.5 channels were found to recover from inactivation prior to deactivation. In addition, it has also been reported (Cummins et al. (2004) that heterologously expressed (in HEK-293 cells) mutant Nav1.7-encoded channels have prolonged deactivation time constants at the hyperpolarized membrane potentials associated with the activation of I_NaR_. These mutant Nav1.7 channels were also reported to display an increase in the relative amplitude of the non-inactivating Nav current (i.e., I_NaP_). These two observations are clearly consistent with the I_NaR_ gating model proposed here in which Nav channels can recover from conventional, fast inactivation directly into an open/conducting state. It is, however, worth noting that in this report (Cummins et al., 2004), I_NaR_ was never measured directly, instead the membrane was repolarized prior to the completion of I_NaT_ inactivation. As a result, it is not possible to conclude, with certainty, that the mutant Nav1.7 channels were recovering from inactivation into an open/conducting state, or simply displaying prolonged deactivation at hyperpolarized membrane potentials.

In developing the model (**Figure 3A**), we found that two fast-inactivated states, i.e., IF1 and IF2, were required to recapitulate the experimental data. It is certainly possible, however, that IF1 and IF2 are substates of the same channel state. or consistency in the model (with single exponential rate parameters), however, we treated these as two distinct states. We also note here that, initially, the model was developed with two slow-inactivated states (IS, IS2) for symmetry. The simulations revealed, however, that the two slow-inactivated states were redundant and that we were able to fit the experimental data reliably with only one of these slow-inactivated states. It is also important to note that we appreciate that our model topology and rate constants, while reliably recapitulating the time- and voltage-dependent properties of the currents determined experimentally, are not necessarily “unique” or potentially the “most simple”.

### Molecular determinants of I_NaR_

Soon after the discovery of I_NaR_, it was reported that the targeted deletion of *Scn8a* (which encodes the Nav1.6 α subunit) in mouse cerebellar Purkinje neurons resulted in a 90% reduction in the amplitude of I_NaR_, suggesting that, of the α subunits (Nav1.1 and Nav1.6) expressed in Purkinje neurons (Vega-Saenz de Miera et al., 1997; Xiao et al., 2013), Nav channels formed by Nav1.6 are the major contributors to I_NaR_ (Raman et al., 1997). However, it was later reported that the targeted deletion of *Scn1a* (which encodes the Nav1.1 α subunit) also results in ∼65% reduction in I_NaR_, suggesting that multiple Nav α subunits contribute to I_NaR_ in cerebellar Purkinje neurons and, in addition, that the contributions are non-linear. To date, all but one (Nav1.3) of the nine Nav channel α subunits have been shown to mediate, or can be induced to mediate, I_NaR_ in native or heterologous cells (Lewis and Raman, 2014, Jarecki et al., 2010, Do and Bean, 2004, Tan et al., 2014). It is unclear whether Nav1.3-encoded Nav channels also generate I_NaR_, as it appears that voltage-clamp studies focused on exploring this possibility have not been conducted to date.

In the model proposed here, the amplitude of I_NaR_ is dependent on, and ultimately regulated by, two factors: the proportion of Nav channels that are non-inactivating at hyperpolarized voltages, i.e., the number of channels that recover from inactivation into an open state on repolarization; and, the proportion of these non-inactivating channels that accumulate into a slow-inactivated state. In the simulations presented in **Figure 9**, we show that by promoting the occupancy of the IS (slow-inactivated) state, we can recapitulate the experimental observation that I_NaR_ amplitudes are reduced in *Scn4b^-/^*, compared with wild type, cerebellar Purkinje neurons, suggesting the possibility that Navβ4 expression regulates I_NaR_ by influencing slow-inactivation. The hypothesis that I_NaR_ is directly affected by Nav channel slow inactivation is also consistent with previous work (Hampl et al., 2016), showing that mutations in the Nav1.6 and Nav1.7 α subunits that enhance or inhibit slow inactivation, also result in reduced or increased, respectively, I_NaR_ amplitudes. Interestingly, it has also been reported that mutation of the IFM motif (to QQQ) in Nav1.4, in addition to completely eliminating fast inactivation, increases the proportion of channels that enter the slow inactivated state (Featherstone et al., 1996), suggesting that the model developed here (**Figure 3A**) in which the fast and slow components of inactivation are exclusive, may be applicable to diverse Nav channels in different cell types and encoded by different α subunits.

There are a number of additional factors, intrinsic and extrinsic, to Nav α subunits, as well as additional pre- and post-translational mechanisms, that regulate persistent Nav currents (Aman et al., 2009, Lin and Baines, 2015, Hammarstrom and Gage, 1998, Paul et al., 2016) and slow-inactivation (Chen et al., 2008, Silva, 2014, Chen et al., 2006, Carr et al., 2003). In the model developed and presented here, the combined effects of these factors/mechanisms will regulate/modulate the amplitudes and the time- and voltage-dependent properties of I_NaR_. In this context, it is interesting to note that the experiments presented in **Figure 6** indicate that the Nav channels underlying I_NaR_, compared to the sum of the Nav channels underlying I_NaT_, are differentially sensitive to slow inactivation. This observation clearly suggests that there is functional (and perhaps molecular) heterogeneity in the population of Nav channels underlying the Nav currents in mouse cerebellar Purkinje neurons, with channels underlying I_NaR_ having distinct properties from the Nav channels underlying I_NaT_.

### Functional Implications

Based on the experimental data and the computational modeling presented here, we propose a novel, blocking particle-independent, gating mechanism for the generation of I_NaR_ that involves two, kinetically-distinct inactivation pathways. The modeling results suggest that two parameters are critical in determining the magnitude and the time- and voltage-dependent properties of I_NaR_: (1) the relative amplitude of the persistent Nav current, I_NaP_, component; and, (2) the proportion of the persistent Nav channels (channels that fail to undergo fast inactivation) that undergo slow inactivation. Interestingly, I_NaR_ has now been identified in over 20 types of neurons, many of which do not display the high rates of repetitive firing that are characteristic of cerebellar Purkinje neurons, suggesting that the role(s) of I_NaR_ in the regulation of membrane excitability are diverse, and likely distinct, in different neuronal cell types (Lewis and Raman, 2014). Recent studies conducted on serotonergic raphe neurons, for example, suggest that the accumulation of Nav channels in a slow-inactivated state functions as a homeostatic brake on repetitive firing (Navarro et al., 2020). In addition, I_NaR_ has been implicated in several inherited and acquired neurological diseases (Lewis and Raman, 2014), including paroxysmal extreme pain disorder, paramyotonia congenita (Jarecki et al., 2010) and chemotherapy-induced neuropathy (Lewis and Raman, 2014, Sittl et al., 2012), as well as epilepsy (Hargus et al., 2013). Given the implications of these findings, it will be of considerable interest to detail the properties of I_NaR_ in different types of neurons and to explore directly the hypothesis that the rate of decay of I_NaR_ on membrane repolarization plays a role in determining how much persistent sodium current is available to contribute to repetitive firing, as well as to define the molecular mechanisms that control I_NaR_ amplitudes, kinetics and functioning in diverse neuronal cell types.

## ACKNOWLEDGMENTS

The authors thank Mr. Richard Wilson for expert technical assistance. The financial support provided by the NIH (R01 NS065761 to JMN, R01 HL136553 to JRS, and F32 NS090765 to JLR) is also gratefully acknowledged; JDM was supported by a NIH institutional training grant (T32 HL007081) and a grant from the Foundation for Barnes Jewish Hospital. The authors declare no competing financial interests.

## AUTHOR CONTRIBUTIONS

All authors contributed to the planning and design of experiments. JLR conducted voltage-clamp experiments. JLR, JMN and JDM analyzed voltage-clamp data. JDM and DB developed Markov models and conducted simulations. All authors contributed to the writing/editing of the manuscript and Figures.

## FIGURE LEGENDS

**Supplemental Figure 1.**
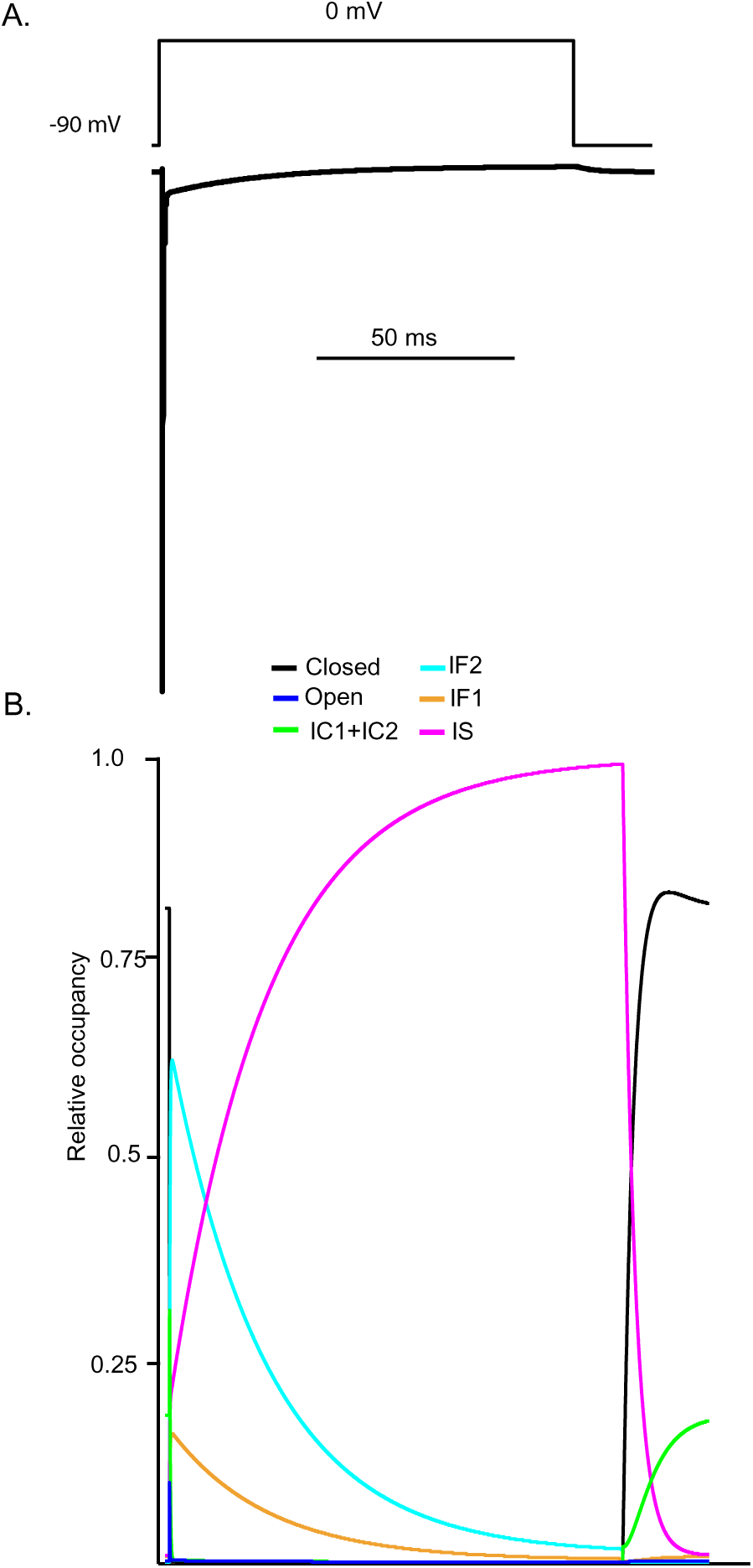
Simulations reveal that, similar to hyperpolarizing voltage steps, prolonged depolarization results in the accumulation of Nav channels in the IS state. Using the novel Markov model developed here (**Figure 3A**), a prolonged depolarizing voltage step to 0 mV produces a rapidly activating Nav current with two kinetically distinct inactivating components (A). The gating state occupancy plot (B) reveals that Nav channels transition first into the IF2 state (*aqua*) and subsequently into the IS state (*orange*).

**Supplemental Figure 2.**
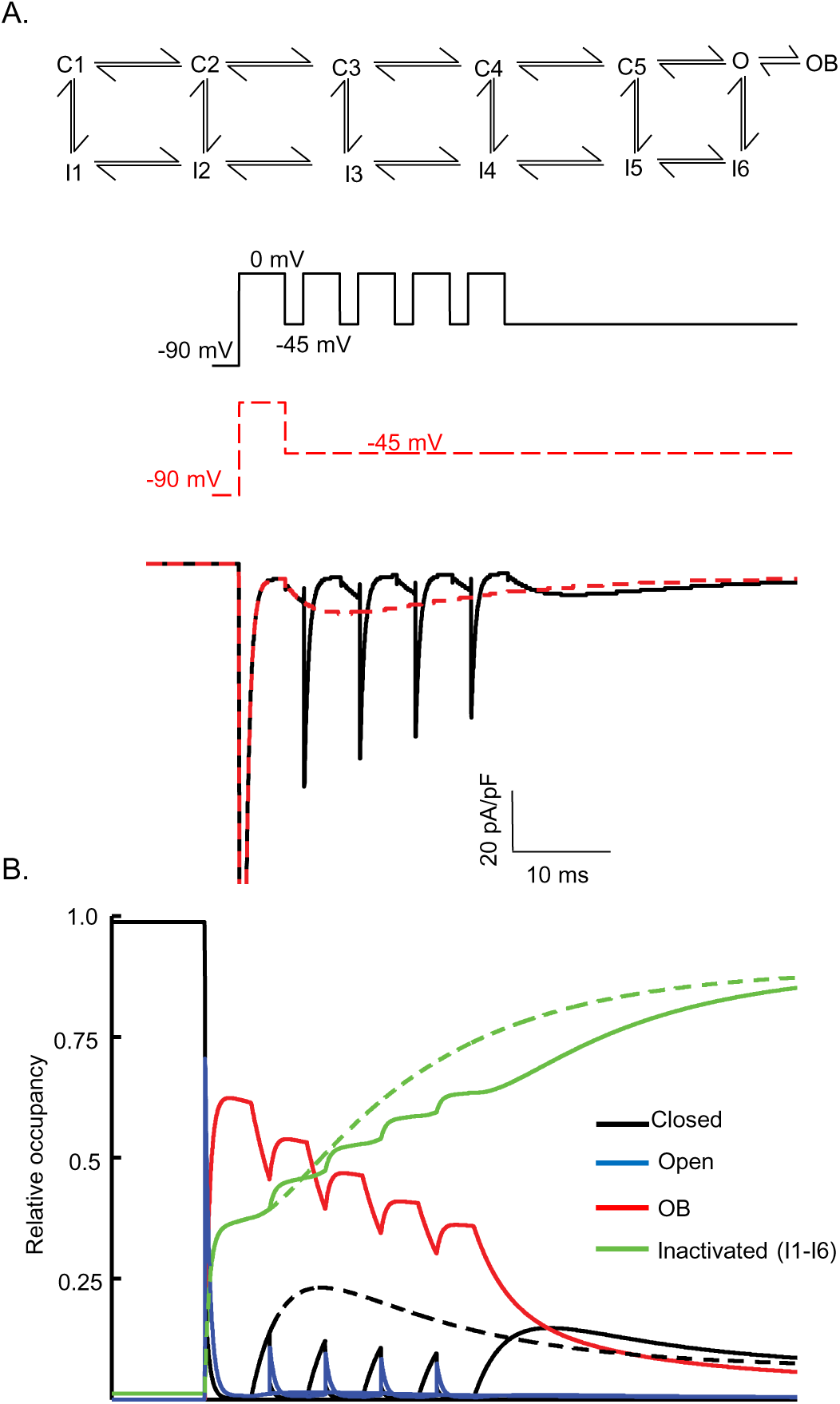
Gating state occupancies/transitions produced in the Raman Bean model (with the voltage-clamp protocols used in Figure 8). The Raman Bean model (presented in **Figure 7A**) does not reproduce the Nav currents recorded from a Purkinje neuron that is depolarized (to 0 mV) and repolarized (to - 45 mV) repeatedly (see: **Figure 8A**); the voltage-clamp protocol and the simulated currents produced are shown in (A) as solid *black* lines. Nav current waveforms, simulated using the Raman Bean model, in response to a sustained hyperpolarization to -45 mV following a brief (5 ms) depolarizing voltage step to 0 mV are also shown; the voltage-clamp protocol and the simulated currents are shown in (A) as dashed *red* lines. The gating state occupancy plot (B), presented on the same time scale as the voltage-clamp records, is shown; colors denote the various kinetic states as indicated. The dashed lines are the gating state transitions associated with the dashed (*red*) current records in (A), and the solid lines are the gating state transitions associated with the solid (*black*) current records in (A).

